# Insight into Cross-Amyloid Interactions and Morphologies: Molecular Dynamics Simulations of Model Peptide Fragments of Amyloid-β (Aβ_16-22_) and Islet Amyloid Polypeptide (IAPP_20-29_)

**DOI:** 10.1101/2021.09.26.461861

**Authors:** N. Cramer, G. Kawecki, K. M. King, D. R. Bevan, A.M. Brown

## Abstract

Amyloid-beta (Aβ) and islet amyloid polypeptide (IAPP) are small peptides, classified as amyloids, that have the potential to self-assemble and form cytotoxic species, such as small soluble oligomers and large insoluble fibrils. The formation of Aβ aggregates facilitates the progression of Alzheimer’s disease (AD), while IAPP aggregates induce pancreatic β-cell apoptosis, leading to exacerbation of Type 2 diabetes (T2D). Cross-amyloid interactions between Aβ and IAPP have been described both *in vivo* and *in vitro*, implying the role of Aβ or IAPP as modulators of cytotoxic self-aggregation of each peptide, and suggesting that Aβ-IAPP interactions are a potential molecular link between AD and T2D. Using molecular dynamics simulations, “hot spot” regions of the two peptides were studied to understand the formation of hexamers in a heterogenous and homogenous peptide-containing environment. Systems of only Aβ_(16-22)_ peptides formed antiparallel, β-barrel-like structures, while systems of only IAPP_(20-29)_ peptides formed stacked, parallel beta strands and had relatively unstable aggregation structures after 2 μs of simulation time. Systems containing both Aβ and IAPP (1:1 ratio) hexamers showed antiparallel, β-barrel-like structures, with an interdigitated arrangement of Aβ_(16-22)_ and IAPP_(20-29)_. These β-barrel structures have features of cytotoxic amyloid species identified in previous literature. Ultimately, this work seeks to provide atomistic insight into both the mechanism behind cross-amyloid interactions and structural morphologies of these toxic amyloid species.

**Statement of Significance:** Molecular knowledge, biophysical characterization, structural morphologies, and formation pathways of amyloid oligomers - specifically low-molecular weight, cross-amyloid oligomers - remain preliminary and undefined. Characterizing interactions between homogenous and heterogenous amyloid oligomers is of great interest given that certain oligomer morphologies contribute to cytotoxicity, eventually resulting in comorbid diseases such as Alzheimer’s disease (AD) and Type 2 Diabetes Mellitus (T2DM). Utilizing model systems (e.g., fragments of full-length peptides) and molecular dynamics (MD) simulations to probe the biophysical underpinnings of cross-amyloid oligomer structures is the first step in understanding the dynamics, stability, and potential modes of cytotoxicity of these species, providing important insights into targetable biomolecular structures.

## Introduction

The accumulation of amyloidogenic peptides into morphologically variable aggregates and highly-ordered plaques are hallmarks of amyloid diseases^1–3^. Amyloid diseases constitute a variety of degenerative disorders, including Alzheimer’s Disease (AD) and Type 2 Diabetes (T2D). In 2020, T2D and AD affected an estimated 6 million^*4*^ and 30 million^*5*^ Americans, and are the sixth and seventh leading causes of death in the US, respectively^*6*^. AD, the most prevalent form of dementia, is characterized by neuronal cell degradation and death^*7*^. T2D is characterized by insulin insensitivity and impaired pancreatic β-cell function, and is exacerbated by poor diet, lack of exercise, and a host of other genetic and environmental factors^*8, 9*^. While both AD and T2D are characterized as individual pathologies, these diseases are highly comorbid. Evidence implicates AD and T2D in mutual pathogenic pathways via overlapping molecular mechanisms^*10–12*^, and each increases the risk for the other.^*13–15*^. Furthermore, AD pathology alters insulin signaling pathways in the brain^*16*^, suggesting AD is implicated in Type III Diabetes^*17*^.

AD and T2D pathologies are associated with the amyloidogenic peptides amyloid-β (Aβ)^*18–21*^ and the islet amyloid polypeptide (IAPP or amylin)^*22–25*^, respectively. Both Aβ and IAPP are capable of self-assembling^*26–29*^ into cytotoxic species of variable morphologies, specifically fibrils^*30–33*^ and oligomers^*20, 34, 35*^. These oligomers, particularly low-molecular weight oligomers, have been identified as key contributors to the onset of each disease and are capable of cross-amyloid interactions.^*36–38*^ Elucidating the mechanisms by which these amyloids aggregate and their resultant structural morphologies can provide insight into the pathogenic events of each disease, allow assessment of the extent of their comorbidity at the molecular level, and provide insight into therapeutic design to disrupt cross-amyloid formation.

Fibrillar amyloid morphologies generally exhibit a cross-β structure, composed of repeating β-strand monomers arranged in a steric zipper^*39, 40*^. Oligomers formed on-pathway to fibrillar aggregate formation are thought to be the primary cytotoxic species in amyloid diseases^*41, 42*^. These have been found to non-specifically destabilize cellular membranes and cause ionic leakage and impaired Ca^2+^ regulation by the formation of β-barrel structures^*43–45*^. Amyloid β-barrels are thought to be composed of a low number of peptide subunits, each assuming a U-shaped ‘β-strand-turn-β-strand’ motif common to most amyloids, and stacked annularly, parallel, and usually in-register to form the barrel.^*46, 47*^ Cytotoxic β-barrels comprised of out-of-register subunits have also been identified^*48*^.

Common structural features of amyloid oligomers and their mechanisms of membrane perturbation have yet to be described at an atomistic level, likely due to their transient nature and widely varying structural morphologies. Molecular dynamics (MD) simulations are ideal for investigating oligomerization events, allowing us to characterize their formation, structure, and stability. To date, MD simulations have been utilized to describe oligomerization of full-length Aβ^*49, 50*^ and IAPP^*51, 52*^, and to determine which portions of their sequences are necessary and sufficient for inducing aggregation and providing stability. In particular, the core hydrophobic regions, composed of residues 16-22 of Aβ (KLVFFAE) and residues 20-29 of IAPP (SNNFGAILSS), have been shown to exhibit similar aggregation potential as full-length peptides^*53, 54*^, and thus are of primary interest as model peptide fragments for studying aggregation.

Previous work characterizing fragments of Aβ and IAPP has provided foundational evidence for modeling these particular fragments. Residues 20-29 of IAPP (IAPP^(20-29)^) form the spine of IAPP fibrils under physiological conditions^*40*^, indicating the ability of this fragment to participate in β-strand formation and its importance for aggregate stability. Furthermore, the primary sequence of this fragment is poorly conserved in IAPP analogues from other species that do not exhibit amyloidogenic properties,^*55*^ indicating that this fragment is influential in the human IAPP aggregation pathway. Likewise, residues 16-22 of Aβ (Aβ^(16-22)^) include the central hydrophobic core of full-length Aβ, which is capable of forming aggregates itself^*56*^ and is essential for the aggregation of the full-length peptide.^*57*^ Furthermore, residues 16-22 of Aβ exist in a natively helical state; a conformational transition of this region to a coiled state capable of integrating into β-strands is considered a major rate-limiting step in Aβ aggregation^*36*^, and interactions between IAPP and this region of Aβ may facilitate this conformational change and subsequently promote co-aggregation.^*36, 58*^ Full-length Aβ and IAPP share approximately 25% sequence identity and 50% sequence similarity, with the greatest conservation occurring within these “hot spot” regions^*59*^; as such, these fragments likely play a central role in cross-amyloid aggregation.

Here, we utilized MD simulations to observe Aβ_(16-22)_ and IAPP_(20-29)_ hexamer formation, using both homogeneous and mixed systems, to provide detailed insight into the structural transitions and architecture of these amyloid fragments. This work describes the underlying structural organization and inter-peptide interactions that these model peptides sample in route to an organized, β-strand rich oligomer structure.

## Methods

The MD simulation package GROMACS (version 2018.1) was used to simulate hexamer formation for all systems using the GROMOS96 53a6 force field.^*60*^ A total of 12 systems (4 replicates of each system type: Aβ_(16-22)_ only, IAPP_(20-29)_ only, or Aβ_(16-22)_+IAPP_(20-29)_ in a 1:1 ratio) were first created by the random placement of 6 copies of peptide fragments in a 13×13×13 nm cubic box. A schematic representation of system design is shown in Figure 1. Peptide fragments of Aβ_(16-22)_ were extracted from full-length Aβ_(1-42)_ (PDB: 1IYT)^*61*^, and fragments of IAPP_(20-29)_ from full-length IAPP_(1-37)_ (PDB: 2L86).^*62*^ All peptide fragments were capped by acetylation at the N-terminus and amidation at the C-terminus. A minimum inter-peptide starting distance of >1.4 nm between the closest two atoms of each peptide fragment was utilized to be beyond the nonbonded cutoff for van der Waals interactions at the onset of the simulation. Systems were then solvated with simple point charge (SPC) water^*63*^ and ionized with 0.150 mM NaCl and counter-ions to preserve a net neutral system.

**Figure 1.**
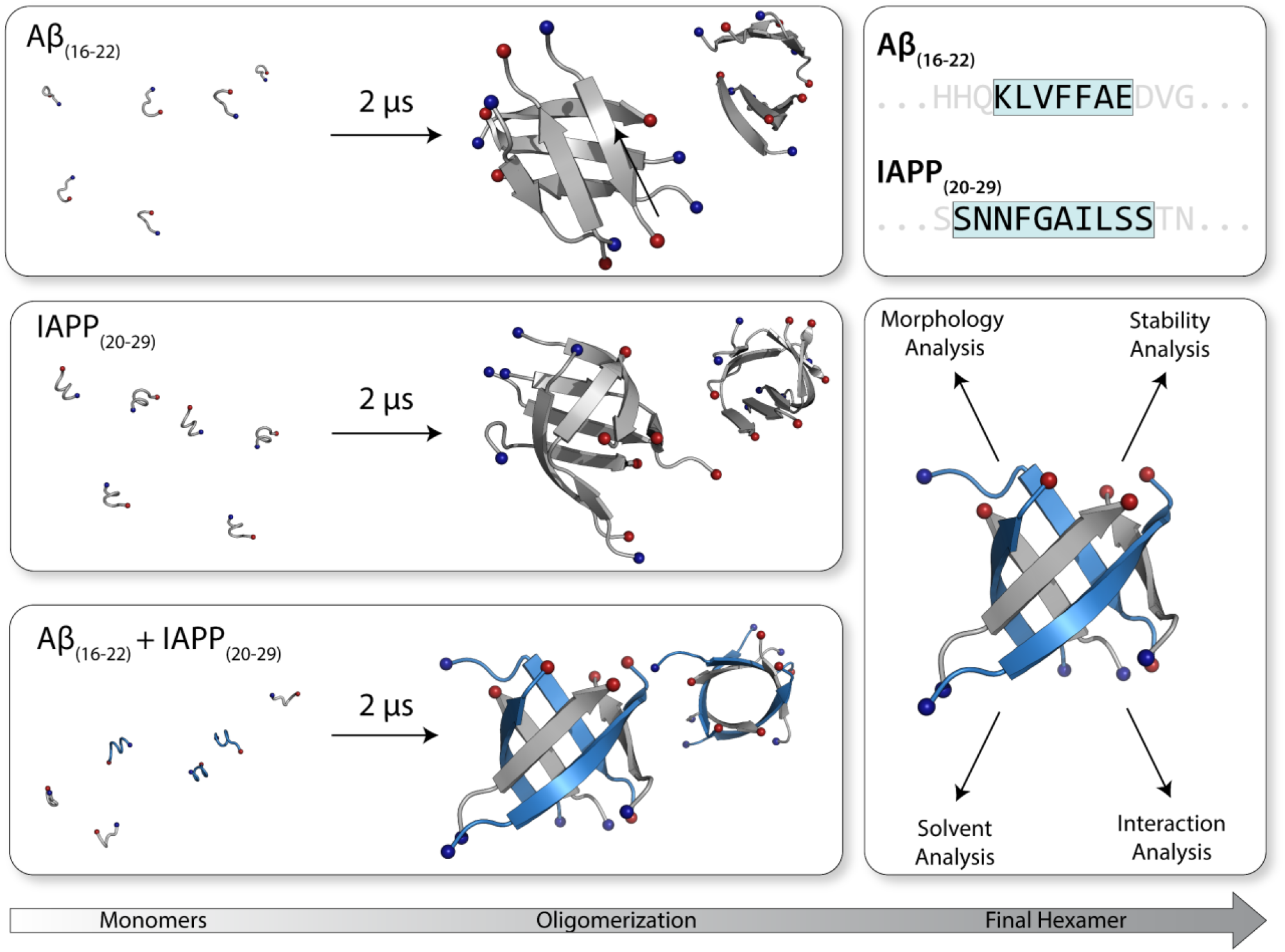
Schematic overview of simulation progression and analysis. Four replicate trajectories per hexamer type were simulated for 2 µs each, providing a cumulative sampling time of 8 µs per hexamer type (Aβ_(16-22)_, IAPP_(20-29)_, and Aβ_(16-22)_ + IAPP_(20-29)_) (Left). Primary sequence of Aβ_(16-22)_ and IAPP_(20-29)_ fragments simulated, highlighted in blue. Overview of analyses performed on final hexamer structures obtained in the last 500 ns of simulation (Right).

The steepest-descent method was employed for energy minimization, after which all systems were subjected to an equilibration by a canonical (NVT) ensemble to maintain temperature of the system at 310K using the Berendsen weak coupling method^*64*^, for 100 ps. Each replicate system utilized different, randomly selected starting velocities assigned at the onset of NVT. An isothermal-isobaric (NPT) ensemble was then employed for a subsequent 100 ps to equilibrate the systems to a pressure of 1 bar using the V-rescale modified Berendsen temperature coupling method^*64, 65*^ and Parrinello-Rahman barostat.^*66, 67*^ Positional restraints were imposed on peptide atoms during equilibration and released afterward. Bond lengths were constrained using Parallel Linear Constraint Solver (P-LINCS)^*68*^ with an integration time step of 2 fs. The smooth Particle-Mesh Ewald method (PME)^*69, 70*^ was used for calculating long-range electrostatics using cubic interpolation. A Fourier grid spacing of 0.16 nm, three-dimensional periodic boundary conditions, and a real-space coulomb cutoff of 1.4 nm were also utilized. Input parameter files, coordinate files, and scripts utilized in this study are located on our Open Science Framework page (https://osf.io/82n73/).

MD production was performed for 2 µs for all replicates in each molecular system for a total simulation time of 24 µs. Root-mean-square-deviation (RMSD) analysis was used to determine simulation convergence for each system; subsequent, aggregate analysis was then conducted over the last 500 ns of each simulation to ensure the characterization of stable hexamers and provide adequate sampling time (6 µs) **(Figures S1-S3)**. RMSD-based clustering of backbone atoms was conducted based on an RMSD cutoff of 0.3 nm, originally based on the work of Darua et al.^*71*^ but expanded to amyloid oligomers in the work of Brown et al.^*72*^, to identify dominant hexamer morphologies for every 100 ns block and for the last 500 ns of each replicate system. Clustered structures were then visualized and rendered in PyMOL (version 2.2.0).^*73*^ Secondary structure was assessed using the Dictionary of Secondary Structure Program (DSSP).^*74*^ Hexameric shape was assessed by calculating eccentricity (*e*) from the three moments of inertia (I_1_, I_2_, and I_3_) and semiaxes *a, b*, and *c*, as previously described.^*72, 75, 76*^ Other analysis, including hydrogen bonding, radius of gyration, and residue-residue contacts were performed using GROMACS analysis tools and in-house scripts. All averages presented were calculated over four replicates for each simulation system and are presented with the corresponding standard deviation. A two-tailed t-test was used for statistical analysis, with statistical significance determined as p < 0.05.

## Results and Discussion

Detailed descriptions of amyloid morphologies and stepwise aggregation events are necessary for understanding and preventing the progression of amyloid diseases like AD and T2D. However, the stochastic nature of these events makes atomistic characterization of aggregate structure particularly difficult. Thus, MD simulations are a valuable tool for providing foundational insight into these mechanisms and identifying potentially targetable structures or co-interaction capabilities. Here, we utilized united-atom MD simulations to model the aggregation of Aβ_(16-22)_ and IAPP_(20-29)_ in both homogenous and heterogenous (mixed) systems. The selected fragments constitute the regions driving oligomerization of Aβ and IAPP, and thus serve as a computationally efficient model system that can be extrapolated to the full-length peptides. Through this work, we provide detailed morphological descriptions of co-aggregated Aβ_(16-22)_ and IAPP_(20-29)_ hexamers and their comparision to homogenous systems, as well as the use of these short peptide fragments for simulating full-length amyloid aggregation events and morphologies.

### Identifying morphologies of homogenous and heterogenous systems of Aβ_(16-22)_ and IAPP_(20-29)_

Adoption of β-strand content is indicative of amyloid aggregation pathways.^*77, 78*^ As such, monitoring the progression of secondary structure allows us to identify canonical on-pathway aggregation. To properly model secondary structure of Aβ _(16-22)_ and IAPP_(20-29)_, force field selection is critical and must be taken into account in system construction and in analysis outcomes. The GROMOS96 53a6 forcefield has been shown to model β-strand formation of Aβ in agreement with experimental data, with other force fields tending to over-stabilize α-helices^71^. For the purposes of this work and based on our previous force field comparison study, the GROMOS96 53a6 was chosen for modelling Aβ and IAPP fragments. A recent review highlights that there are three major problems to consider in simulating amyloid aggregation, including insufficiently accurate force fields, protein concentration, and time-scale limits.^*79*^ More work is necessary to further refine force fields for both the field of IDPs, for model systems, and for aggregation processes, with this work contributing how one force field models similarities and differences between model systems of homogenous and mixed amyloid formation. This work focuses on homogenous and mixed amyloid structure formation on that facet and does not seek to be a comparision between force fields. Furthermore, recent work from discrete MD (DMD) simulations of full-length Aβ_42_ hexamer formation indicate a β-barrel formation in line with the results discussed below, as well as *in vivo* experimental validation, that highlighted the formation the β-barrel oligomers of Aβ_42_^*80*^. To describe aggregate structure with respect to secondary structure content, secondary structure over time for each system was assigned using the DSSP algorithm^*81*^. Aβ control hexamers showed a rapid timewise transition from 100% random coil at MD onset to 51% ± 4 β-strand structure over the last 500 ns of simulation **(Figure 2A, Table 1)**. Likewise, IAPP control hexamers showed a similarly rapid progression from predominantly random coil at the onset of MD, stabilizing at 48% ± 8 β-strand structures in the last 500 ns of simulation **(Figure 2B, Table 1)**. These data match previous analyses of secondary structure content of amyloid oligomers, which adopt β-strand structure on-pathway to aggregation^*82,83*^. Fragments adopting experimentally determined secondary structure content supports the use of GROMOS 53a6 for modelling these fragments.

**Table 1.** Average post-convergence secondary structure percentages and standard deviations by system. Values were obtained by sampling the last 0.5 µs of four replicate trajectories per system for a total sampling time of 2 µs per hexamer type.

| System | Coil | β-Strand | α-Helix |
| --- | --- | --- | --- |
| Aβ Control | $49 \pm 4$ | $51 \pm 4$ | $0 \pm 0$ |
| IAPP Control | $52 \pm 8$ | $48 \pm 8$ | $0 \pm 0$ |
| Aβ+IAPP | $58 \pm 8$ | $42 \pm 8$ | $0 \pm 0$ |

**Figure 2.**
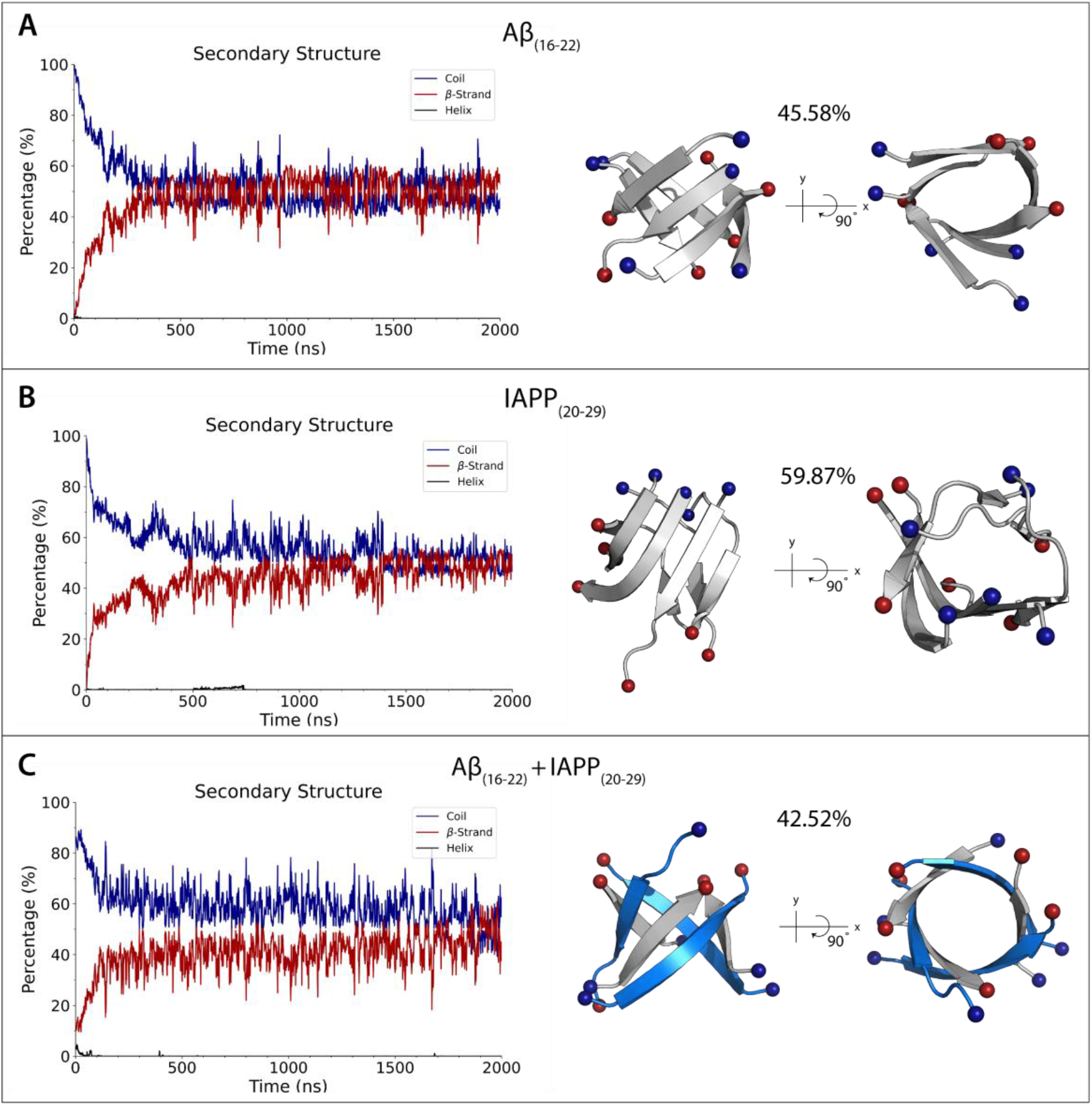
Secondary structure over time and representative structures from RMSD clustering for Aβ_(16-22),_(B) IAPP_(20-29)_, and (C) Aβ_(16-22)_ + IAPP_(20-29)_. The percentage of residues exhibiting each secondary structure type was averaged across four replicate trajectories for each system type, and block averaged every 1 ns are shown. Cluster structures from replicates with the highest cluster percentage and similarity to average secondary structure of all replicates represent morphologies of the last 500 ns of simulation. **(A)** AB_(16-22)_ is colored gray, **(B)** IAPP_(20-29)_ is colored gray **(C)** IAPP_(20-29)_ is colored blue, and AB_(16-22)_ is colored gray. All peptides are shown as cartoon. The N- and C-termini are shown as spheres, and colored blue and red, respectively.

Mixed Aβ+IAPP hexamers exhibited the same rapid progression from predominantly coil to greater β-strand content, reaching 42% ± 8 β-strand content in the last 500 ns, though the percentage is lower than observed for Aβ_(16-22)_ and IAPP_(20-29)_ control systems **(Figure 2C, Table 1)**. This apparent decrease in β-strand content may be attributed to differing length of the two fragment types (7 residues for Aβ_(16-22)_ vs 10 residues for IAPP_(20-29)_). Indeed, structures from RMSD clustering show the terminal ends of the longer IAPP(20-29) fragment adopting random coil secondary structure **(Figure S4)**.

Given the polymorphic nature of amyloid oligomers, the most targetable motifs are likely found in tandem with the most stable or common morphologies. Thus, to characterize dominant morphologies of each hexamer type, clustering analysis was performed over the last 500 ns of simulation **(Figure 2, Figures S5-S7)** and over 100 ns time periods over the 2 µs simulation **(Figures S8-S19)**. Dominant morphologies for Aβ_(16-22)_ replicates over the last 500 ns of simulation included β-barrel-like structures and stacked β-strand pairs consisting of trimer/trimer and tetramer/dimer configurations **(Figure 2A, Figures S5, S8-S11)**. β-strands tend to be antiparallel, although some parallel strands are observed. These structures are in agreement with previous literature; Aβ oligomers favor a variety of comparable structures based on the lack of a single local minima in a free energy landscape, adopting several localized energy basins representing stacked beta strands^*84, 85*^ or barrel-like structures^*86, 87*^.

IAPP_(20-29)_ oligomers instead formed either shorter-length β-strands interspersed by random coil, or a single, tightly curved strands reminiscent of open-conformation β-barrels **(Figure 2B, Figures S6, S12-S15)**. Both parallel and anti-parallel β-strands are observed, although parallel is dominant. Computational work performed by Sun et al.^*50*^ describes a progression of events during IAPP aggregation that may provide a mechanical link between the structures seen here, in which oligomerized peptides initially form short β-strands flanked by random coil, which then elongate and form a single β-strand capable of curling and adopting a barrel-like structure. The apparent frequency of the structures obtained for Aβ_(16-22)_ and IAPP_(20-29)_ in the literature suggest that these structures are representative of on-pathway oligomers.

Mixed hexamers composed of Aβ_(16-22)_ and IAPP_(20-29)_ generally exhibited barrel-like morphologies, with all four systems showing noticeable pore-like architecture **(Figure 2C, Figures S7, S16-S19)**. Typically, mixed Aβ_(16-22)_ and IAPP_(20-29)_ structures displayed an interdigitated arrangement of Aβ_(16-22)_ and IAPP_(20-29)_ peptides, usually arranged in parallel β-strands. Similar β-barrel structures have been observed in other computational work studying Aβ-IAPP fragment co-aggregation^*88*^. Furthermore, hydrogen exchange mass spectrometry^*89*^ and NMR^*86*^ experiments with full-length Aβ reveal that β-barrel oligomers are formed on pathway to aggregation, suggesting that the fragment model system is representative of full-length co-aggregation behavior. Interestingly, the β-barrel oligomer formed by the amyloidogenic αB crystallin peptide, named cylindrin, has been found to exhibit cytotoxicity^*90*^. Thus, it is possible that co-aggregated IAPP and Aβ exert cytotoxicity via β-barrel formation and when in mixed amyloid systems, have a higher propensity for β-barrel formation and more ordered structures than when in homogenous systems.

Determining aggregate structures for basic mechanistic insight into disease progression is dependent on structural stability. As such, we sought to determine the relative stability of each hexamer. To estimate structural stability, cluster structure percentages, solvent-accessible surface area (SASA), and free energy calculations were examined. Clustering percentages indicate the percentage of frames within 0.3 nm of a given structure; as such, higher clustering percentages signify more conformational stability. On average, Aβ_(16-22)_ hexamers were determined to be more stable and unchanging morphologically than both IAPP_(20-29)_ and mixed Aβ_(16-22)_ + IAPP_(20-29)_ hexamers as indicated by clustering percentages calculated from the last 500 ns of each simulation **(Table 2)**. Interestingly, Aβ hexamers exhibited the most favorable total free energy despite having a larger hydrophobic solvent-accessible surface area relative to IAPP and mixed systems **(Table 2)**. This increased stability is likely due to a combination of aromatic-hydrophobic and charged inter-peptide interactions. Aromatic interactions were observed to be high-probability interactions in residue network maps **(Figure 3)**, and cluster structures revealed that solvent-exposed phenylalanine residues exhibited ordered stacking interactions in replicates with less structural polymorphism **(See Figure S20)**. Furthermore, the charged side chains on the terminal residues of Aβ_(16-22)_ likely contribute to structural stability; the average minimum distance between Aβ terminal residues (0.93 ± 0.04 nm) are significantly different from distances between terminal residues IAPP (1.16 ± 0.03 nm) or mixed systems (1.70 ± 0.05 nm), which are polar/polar and charged/polar, respectively **(Table 2)**. This indicates that contacts between Aβ terminal residues are closer; potential salt bridges were observed in replicates 1, 2, and 3 **(Figure S21)**. These interactions between charged terminal residues may account for the increased stability seen in Aβ systems.

**Table 2.**
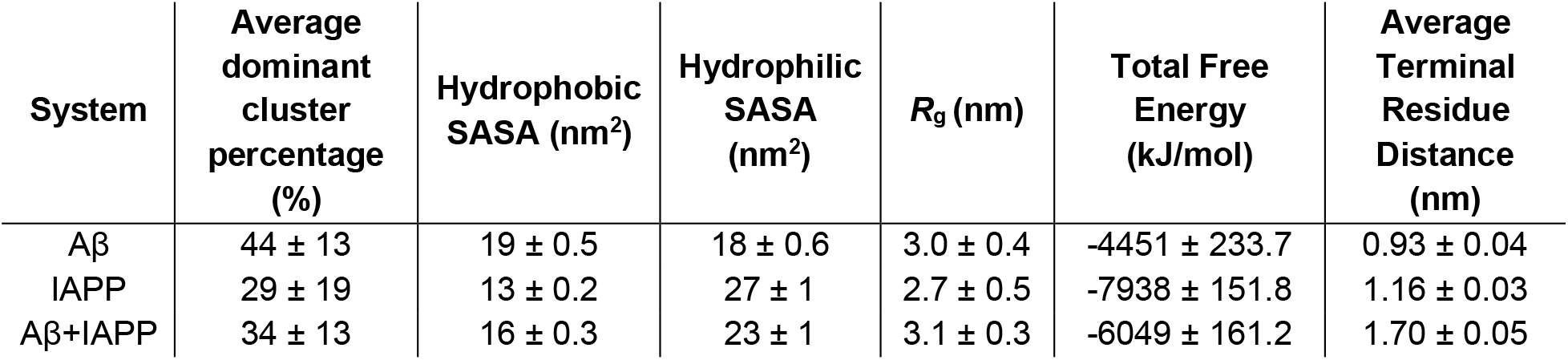
Dominant cluster percentage, solvent accessible surface area (SASA) radius of gyration, free energy averages, and average distance between polar/charged terminal residues. Values were obtained by sampling the last 0.5 µs of four replicate trajectories per system for a total sampling time of 2 µs per hexamer type.

| System | Average dominant cluster percentage (%) | Hydrophobic SASA (nm<sup>2</sup>) | Hydrophilic SASA (nm<sup>2</sup>) | $R_g$ (nm) | Total Free Energy (kJ/mol) | Average Terminal Residue Distance (nm) |
| --- | --- | --- | --- | --- | --- | --- |
| Aβ | 44 ± 13 | 19 ± 0.5 | 18 ± 0.6 | 3.0 ± 0.4 | -4451 ± 233.7 | 0.93 ± 0.04 |
| IAPP | 29 ± 19 | 13 ± 0.2 | 27 ± 1 | 2.7 ± 0.5 | -7938 ± 151.8 | 1.16 ± 0.03 |
| Aβ+IAPP | 34 ± 13 | 16 ± 0.3 | 23 ± 1 | 3.1 ± 0.3 | -6049 ± 161.2 | 1.70 ± 0.05 |

**Figure 3.**
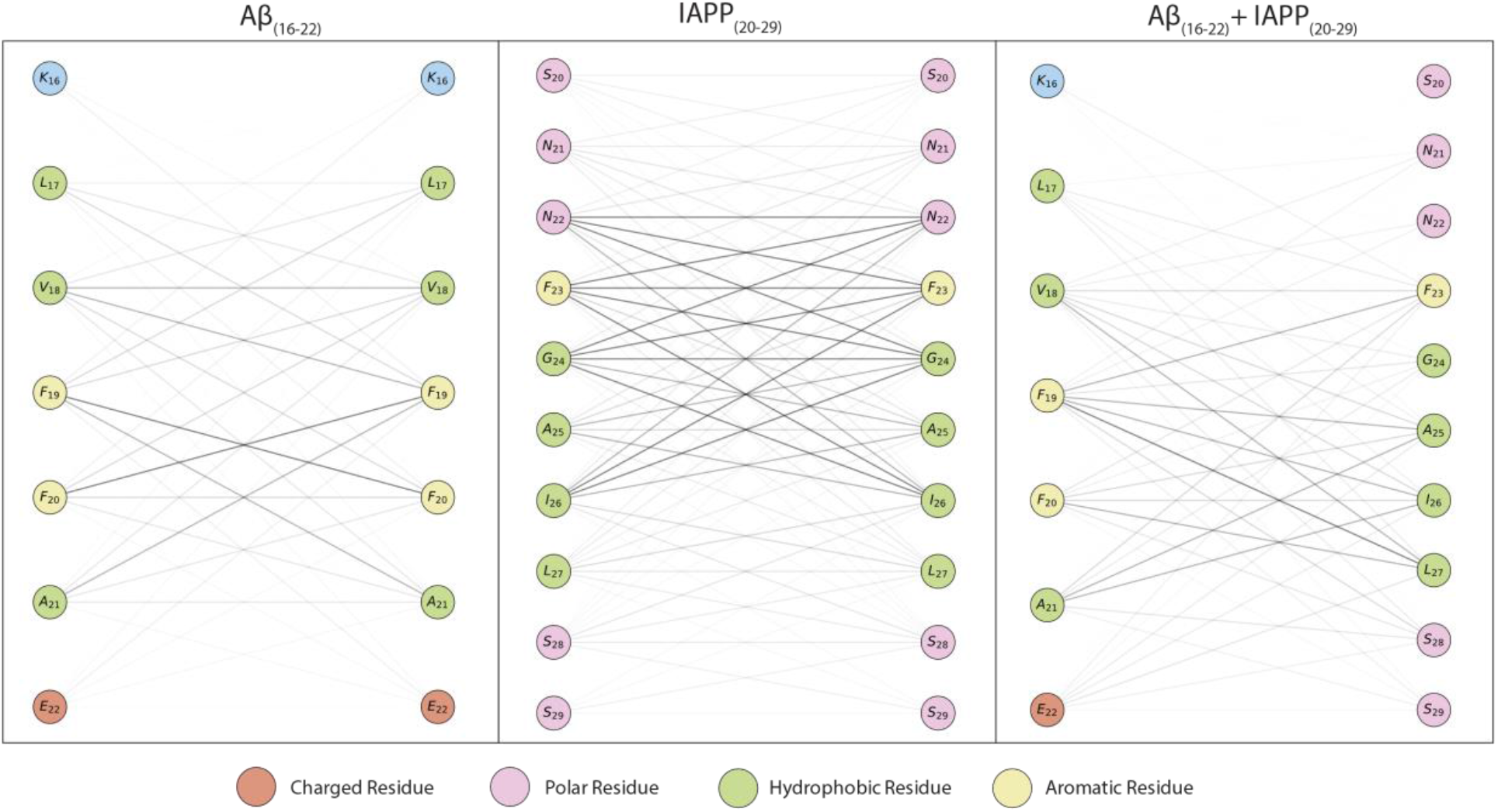
Network maps depicting frequency of interaction in the last 500 ns of simulation between residues for (left) Aβ_(16-22)_, (center) IAPP_(20-29)_, and (right) AB_(16-22)_ + IAPP_(20-29)_. Interaction frequency was normalized by number of interacting atoms. Darkness of lines is proportional to frequency of interaction.

IAPP_(20-29)_ hexamers were the most polymorphic of any of the systems, exhibiting the most numerous cluster structures sampled over the last 500 ns of simulation **(Table 2)**. Furthermore, the free energy of IAPP systems was on average the most favorable with the lowest free energy calculation of the final formed morphologies, with the lowest amount of hydrophobic SASA over the last 500 ns of simulation **(Table 2)**. IAPP, like Aβ, benefits from strong interactions between hydrophobic residues **(Figure 3)**. We predict that IAPP systems were more energetically favorable given the decrease in order and barrel formation observed as compared to Aβ only systems. Interestingly, IAPP is modestly more compact than the Aβ and mixed systems as assessed by radius of gyration, suggesting that hydrophobic packing is strongest in IAPP systems **(Table 2)**. The reduced stability in IAPP systems is likely due to weaker interactions from flanking polar residues relative to Aβ; average minimum distance between the polar termini was longer than distance between charged terminal residues in Aβ **(Table 2)**. The polar termini are thus more flexible, and likely contribute to the structural polymorphism.

Mixed Aβ_(16-22)_ + IAPP_(20-29)_ hexamers mediate characteristics of homogenous Aβ_(16-22)_ and IAPP_(20-29)_ systems in terms of stability based on clustering, free energy, SASA and compactness **(Table 2)**. Like their substituents, mixed systems likely derive stability from strong hydrophobic interactions between the cores of each peptide, where tight hydrophobic packing is observed in network maps **(Figure 3)**. In particular, Aβ Phe residues have high probability of interaction with other Aβ Phe and IAPP Leu residues, packing in the core of the barrel **(Figure S22)**. However, much like IAPP_(20-29)_, the mixed systems exhibited less frequent contacts between flanking polar/charged residues than did Aβ or IAPP **(Table S2)**. This can be attributed to the differing length between the two peptides, often leaving the N-terminal region of IAPP disordered in the simulations. There was a significant increase in free energy favorability of the mixed Aβ_(16-22)_ + IAPP_(20-29)_ compared to homogenous systems of Aβ. While the overall morphology of Aβ_(16-22)_ + IAPP_(20-29)_ systems mimicked Aβ-only systems, and the structural morphology of a β-barrel was more consistent and defined, the energetic favorability for a more ordered structure was observed. This indicated that the presence of both peptides in a mixed system contributed to a more favorable total free energy and ordered structure than individually, which highlights the potentiality of exacerbation of oligomer toxicity in co-amyloid structures.

### Structural implications related to cytotoxicity of cross-amyloid structures

Combating amyloid diseases like AD and T2D also requires translating these differences in oligomeric structure and stability to relative toxicity. Out-of-register β-strand alignment, leaving staggered hydrogen bonds, has been implicated in the formation of toxic, cylindrin-like amyloids^*91*^. Out-of-register β-strands were observed in mixed Aβ_(16-22)_ + IAPP_(20-29)_ replicates. We calculated the average number of hydrogen bonds between residues and water for each system to determine if out-of-register hydrogen bonds are of higher frequency in heterogeneous systems. Hydrogen bonding with water was increased in mixed systems, particularly with respect to N-terminal IAPP_(20-29)_ residues Asn21 and Asn22 **(Figure S23)**. If hydrophobic core contacts are preserved in full-length cross-aggregates, out-of-register hydrogen bonds could potentially be present on the N-terminal end of IAPP. This supports the formation of β-barrels for full length cross-aggregates and may provide insight into their toxicity. Additionally, Aβ Glu22 residues were more likely to have an increase in number of hydrogen bonds with water, in both the mixed and homogenous system.

The formation of several distinct pore-like amyloid oligomers have been reported in the literature. It is hypothesized that these structures may confer toxicity by permeabilizing the membrane to ions, leading to cellular dysfunction. While the β-barrel structures observed appear pore-like in shape, the internal cavity of the barrel is generally packed with hydrophobic side chains. As such, barrel-like structures are accordingly unlikely to accommodate passage of significant quantities of water, corroborated by water shell analysis **(Figure 4, Figures S24-S26)**. Similar hydrophobic packing has been observed in other studies examining the formation of pore-like Aβ structures^*90*^. In general, more apparent barrel-like structures appear to accommodate notably less water in the internal cavity due to increased hydrophobic packing, and therefore have less internal surface area for water contact. However, it is worth mentioning that permeability to solvent and ions may be dependent on oligomer size; Aβ 20-mers appear to have characteristics of ion channels^*92*^. Furthermore, disordered oligomers with increased hydrophobic SASA have been identified as cytotoxic^86, 87^, suggesting that water permeability may be a factor in cytotoxicity.

**Figure 4.**
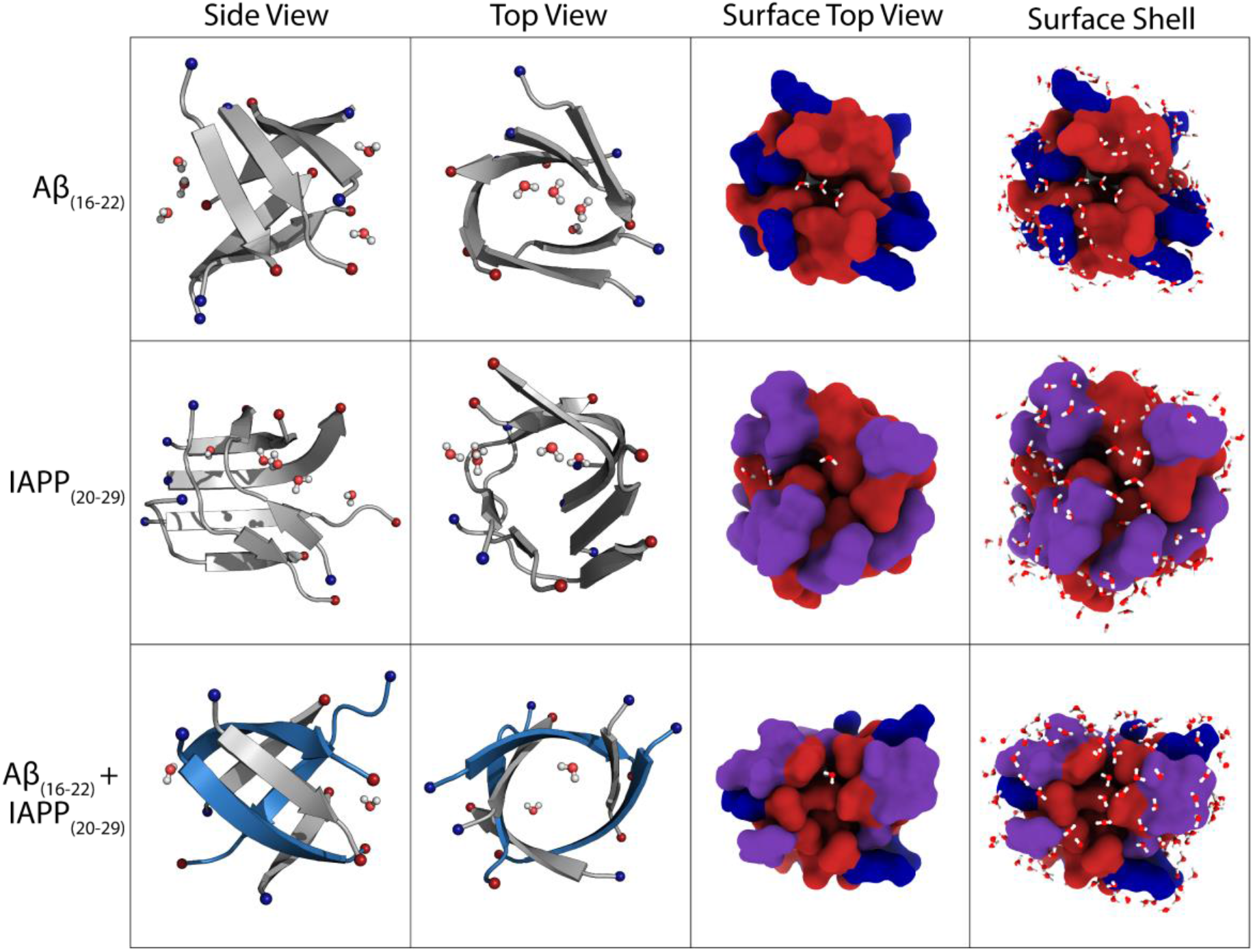
Internal water molecules and water molecules within 3 Å of solvent-exposed residues (surface shell). Each morphology represents the central structure of the dominant cluster obtained from the last 0.5 µs of each system. In the left-most columns, IAPP_(20-29)_ fragments are shown in blue, while Aβ_(16-22)_ fragments are shown in gray. In the rightmost columns, peptides are shown as a surface, and colored by hydrophobicity (charged blue, polar purple, hydrophobic red).

## Conclusions

This work indicates that mixed systems containing both Aβ_(16-22)_ and IAPP_(20-29)_ form β-barrel oligomers, which are potentially cytotoxic and have different structure morphologies and stability when compared to homogenous Aβ_(16-22)_ or IAPP_(20-29)_ hexamers. This difference in homogenous versus mixed amyloid systems is supported by both the clustering analysis and eccentricity analysis, as well as average termini distances. Mixed systems mediate characteristics of both Aβ_(16-22)_ and IAPP_(20-29)_, with the presence of Aβ likely organizing and influencing β-strand content and morphology over IAPP. Mixed systems of Aβ_(16-22)_ and IAPP_(20-29)_ consistently formed more organized, interdigitated β-barrel shaped morphologies than homogenous systems. This work highlights the utilization of short, model fragments to simulate the similarities and differences between homogenous and heterogenous amyloid structures, Future work includes molecular dynamics simulations with full length Aβ and IAPP mixed systems to see co-amyloid behavior is comparable to the results obtained from using fragments as model systems, as well as the impact of these morphologies on membrane perturbation as a metric to assess cytotoxicity.

## Author Contributions

N.C, D.R.B. and A.M.B. designed research protocol. N.C, G.K., and A.M.B. performed simulations. N.C., G.K, K.M.K, and A.M.B analyzed data. N.C., G.K, D.R.B, and A.M.B. wrote the paper.

## Acknowledgments

The authors thank Advanced Research Computing at Virginia Tech for high performance computing resources.

