## Supplemental Information for "Insight into Cross-Amyloid Interactions and Morphologies: Molecular Dynamics Simulations of Model Peptide Fragments of Amyloid-β (Aβ_16-22_) and Islet Amyloid Polypeptide (IAPP_20-29_)"

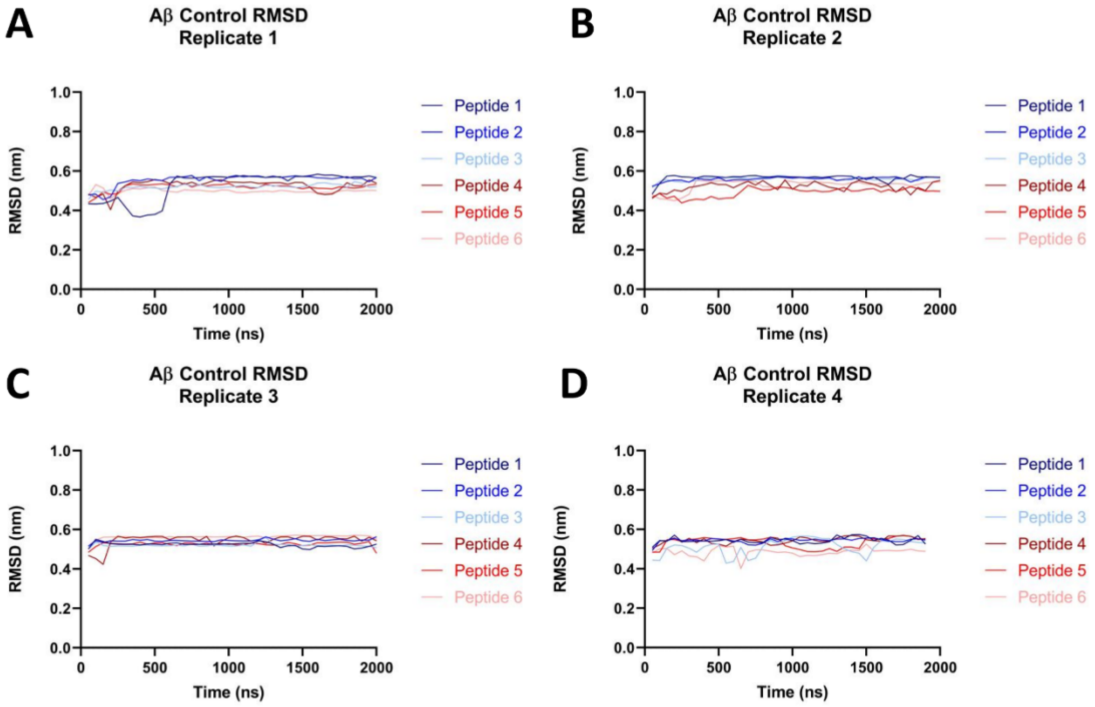


**Figure S1. RMSD of Aβ$_{(16\text{-}22)}$ control backbone atoms.** RMSD values were calculated using the starting positions of the peptides as a reference frame. Values were then block averaged in 50 ns blocks and plotted as a function of time. RMSD of Aβ control systems shows stable hexamer formation between 500 and 1000 ns for most systems. Individual peptide RMSD was calculated and is shown by varying colors of blue and red.

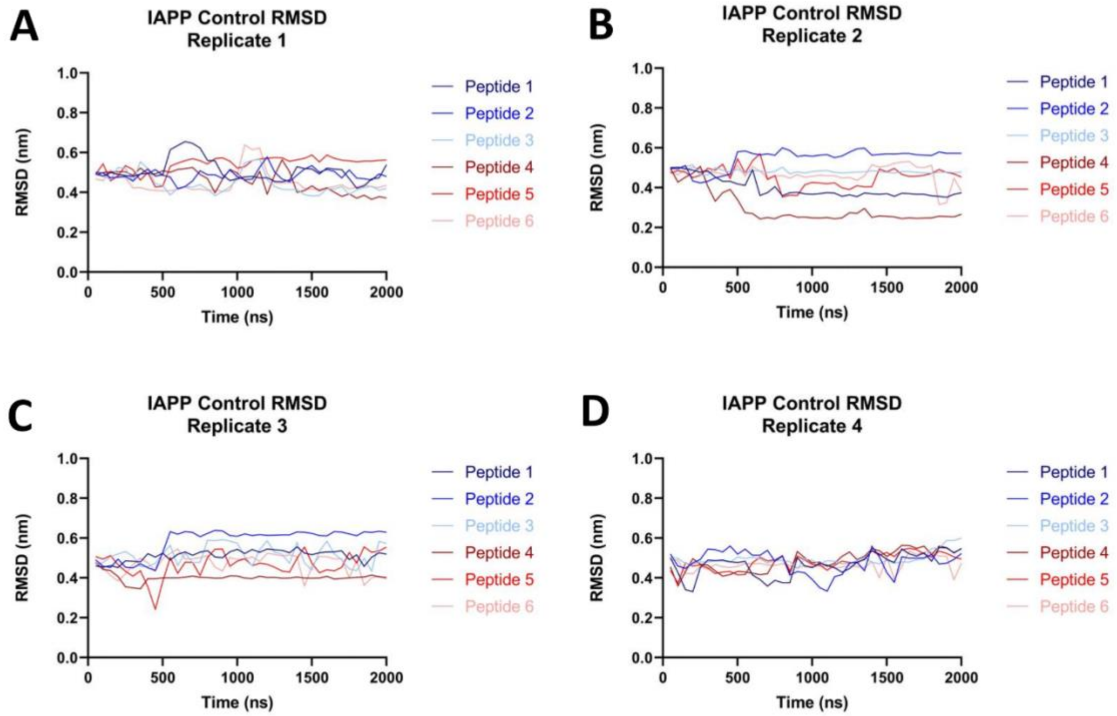


**Figure S2. RMSD of IAPP$_{(20\text{-}29)}$ control backbone atoms.** RMSD values were calculated using the starting positions of the peptides as a reference frame. Values were then block averaged in 50 ns blocks and plotted as a function of time. RMSD of IAPP control systems indicates that stable hexamer formation generally occurred after 1000 ns, indicating a longer aggregation mechanism. The more variable values for each peptide suggest that IAPP control hexamers are less ordered relative to Aβ control hexamers.

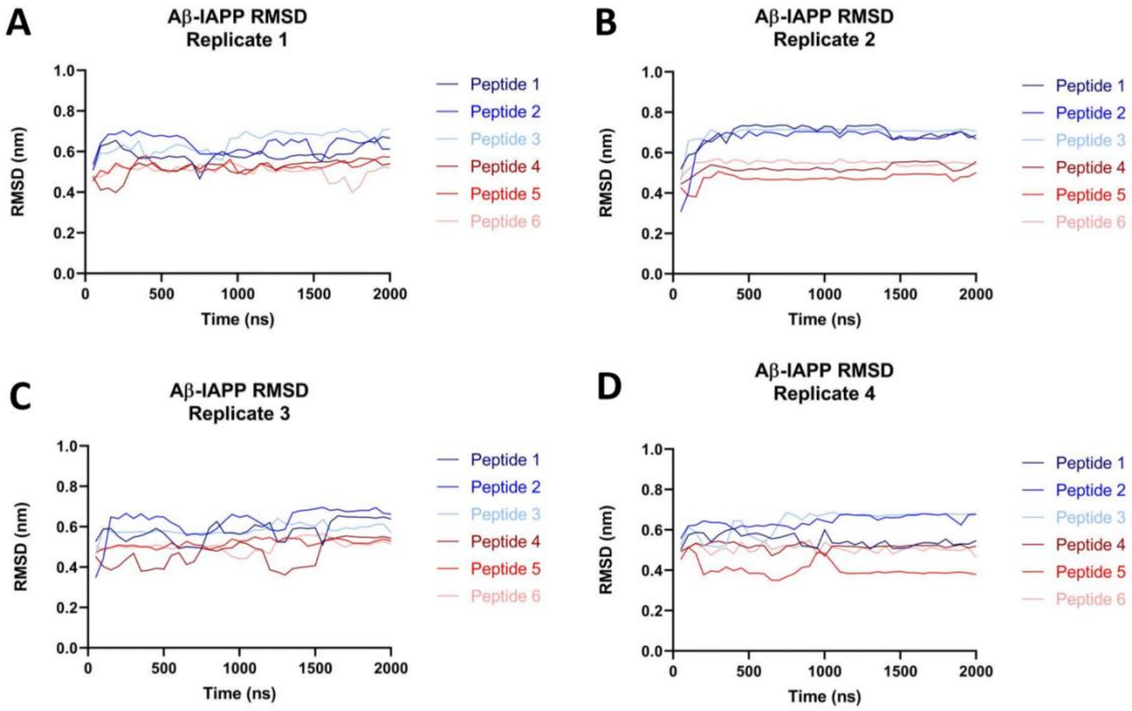


**Figure S3. RMSD of Aβ$_{(16\text{-}22)}$+IAPP$_{(20\text{-}29)}$ backbone atoms.** RMSD values were calculated using the starting positions of the peptides as a reference frame. Values were then block averaged in 50 ns blocks and plotted as a function of time. IAPP monomers are shown in blue, while Aβ monomers are shown in red. RMSD of heterogeneous hexamers resembles the IAPP control hexamers, in that stable hexamer formation generally took longer to occur (post-1000 ns), and hexamers were more variable.

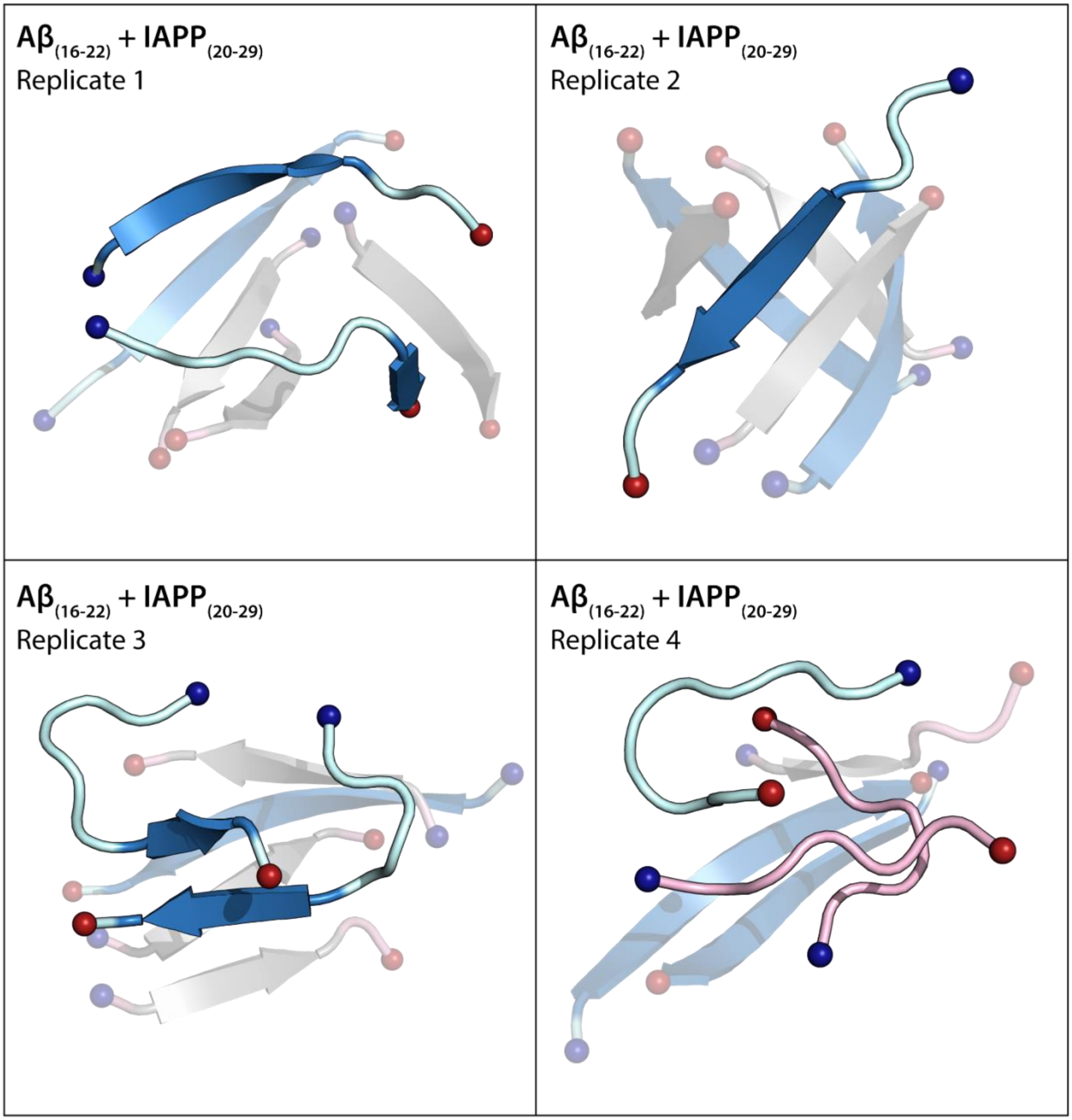


**Figure S4. Random coil adoption by IAPP$_{(20-29)}$ in heterogenous Aβ$_{(16-22)}$ + IAPP$_{(20-29)}$ systems.** Aβ$_{(16-22)}$ β-strand structure is colored gray, and random coil is colored light pink. IAPP$_{(20-29)}$ β-strand structure is colored blue, with random coil structure colored light blue. All peptides shown as cartoon. N- and C- termini are shown as spheres, and colored blue and red, respectively.

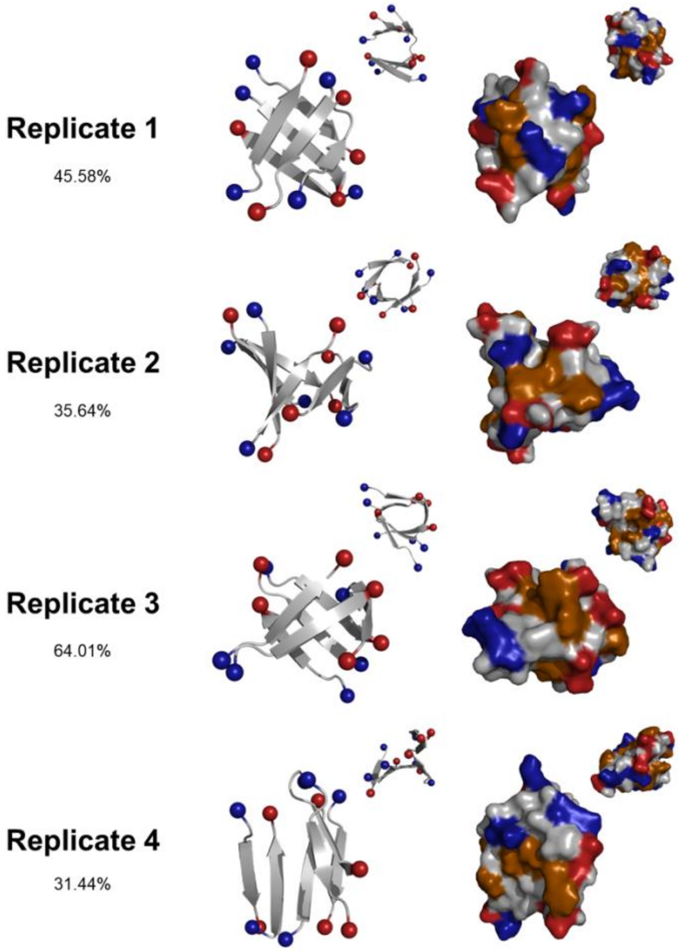


**Figure S5. Dominant morphologies and surface representations of Aβ Control hexamers.** Structures shown are the dominant cluster morphology obtained from backbone RMSD clustering over the final 500 ns of each simulation. Structures are shown as secondary structure and surface. Secondary structure structures are shown as cartoon, colored grey, with N- and C-termini colored blue and red, respectively. Surface representations are colored by the chemical properties of surface-accessible residues (gray: aliphatic, blue: positively charged, red: negatively charged, orange: aromatic, and purple: polar uncharged). Percentages indicate the percentage of frames belonging to the most dominant cluster over the last 500 ns of simulation time.

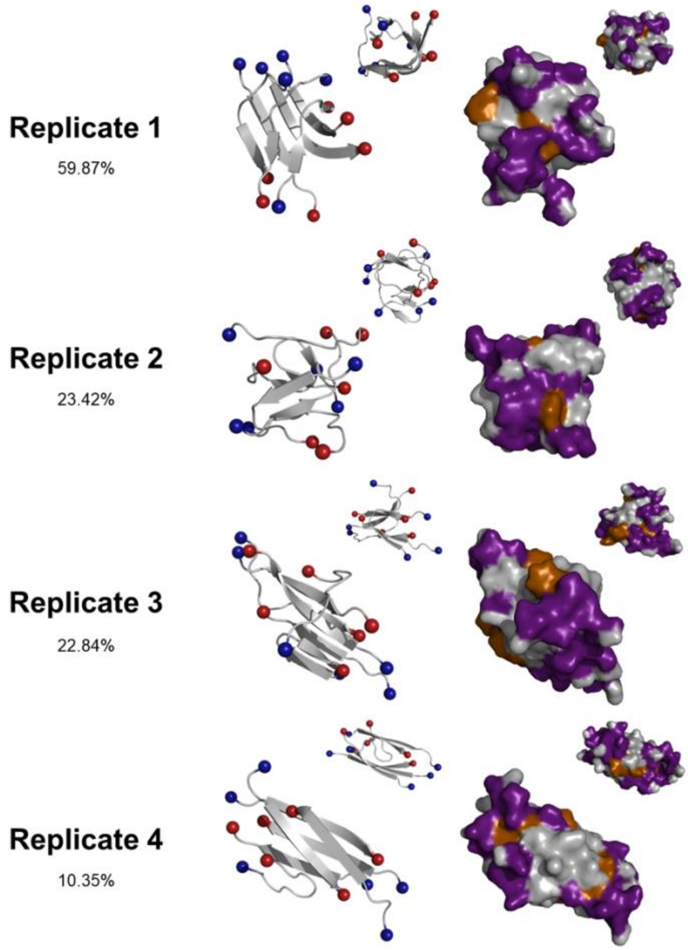


**Figure S6. Dominant morphologies and surface representations of IAPP control hexamers.** Structures shown are the dominant cluster morphology obtained from backbone RMSD clustering over the final 500 ns of each simulation. Structures are shown as secondary structure and surface. Secondary structure structures are shown as cartoon, colored grey, with N- and C-termini colored blue and red, respectively. Surface representations are colored by the chemical properties of surface-accessible residues (gray: aliphatic, blue: positively charged, red: negatively charged, orange: aromatic, and purple: polar uncharged). Percentages indicate the percentage of frames belonging to the most dominant cluster over the last 500 ns of simulation time.

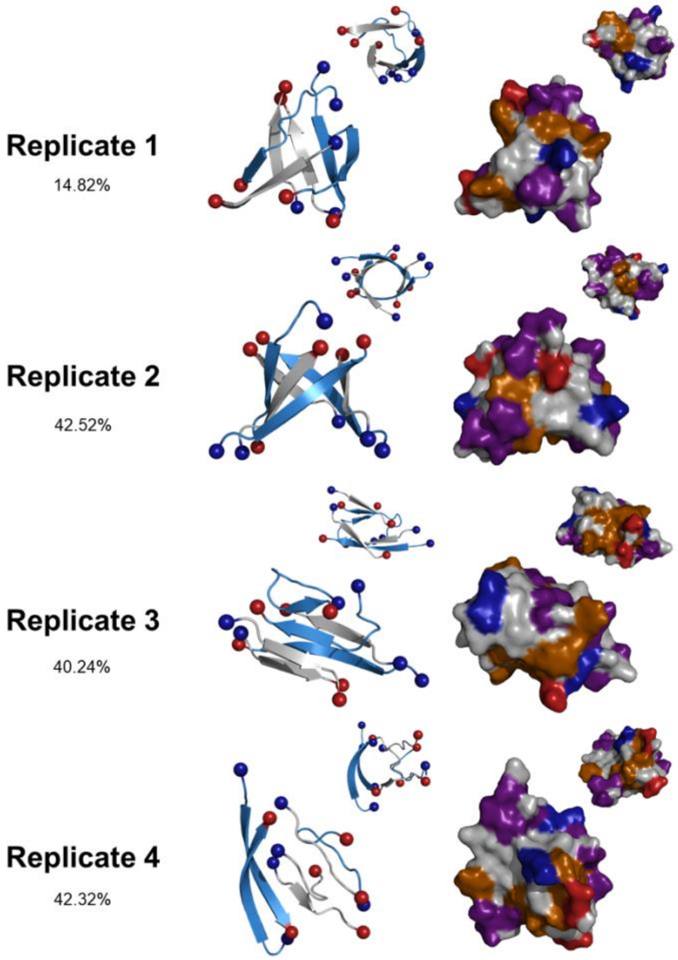


**Figure S7. Dominant morphologies and surface representations of mixed Aβ+IAPP hexamers.** Structures shown are the dominant cluster morphology obtained from backbone RMSD clustering over the final 500 ns of each simulation. Structures are shown as secondary structure and surface. Secondary structure structures are shown as cartoon, with N- and C-termini colored blue and red, respectively. $IAPP_{(20-29)}$ fragments are colored blue, and $A\beta_{(16-22)}$ peptides are colored gray. Surface representations are colored by the chemical properties of surface-accessible residues (gray: aliphatic, blue: positively charged, red: negatively charged, orange: aromatic, and purple: polar uncharged). Percentages indicate the percentage of frames belonging to the most dominant cluster over the last 500 ns of simulation time.

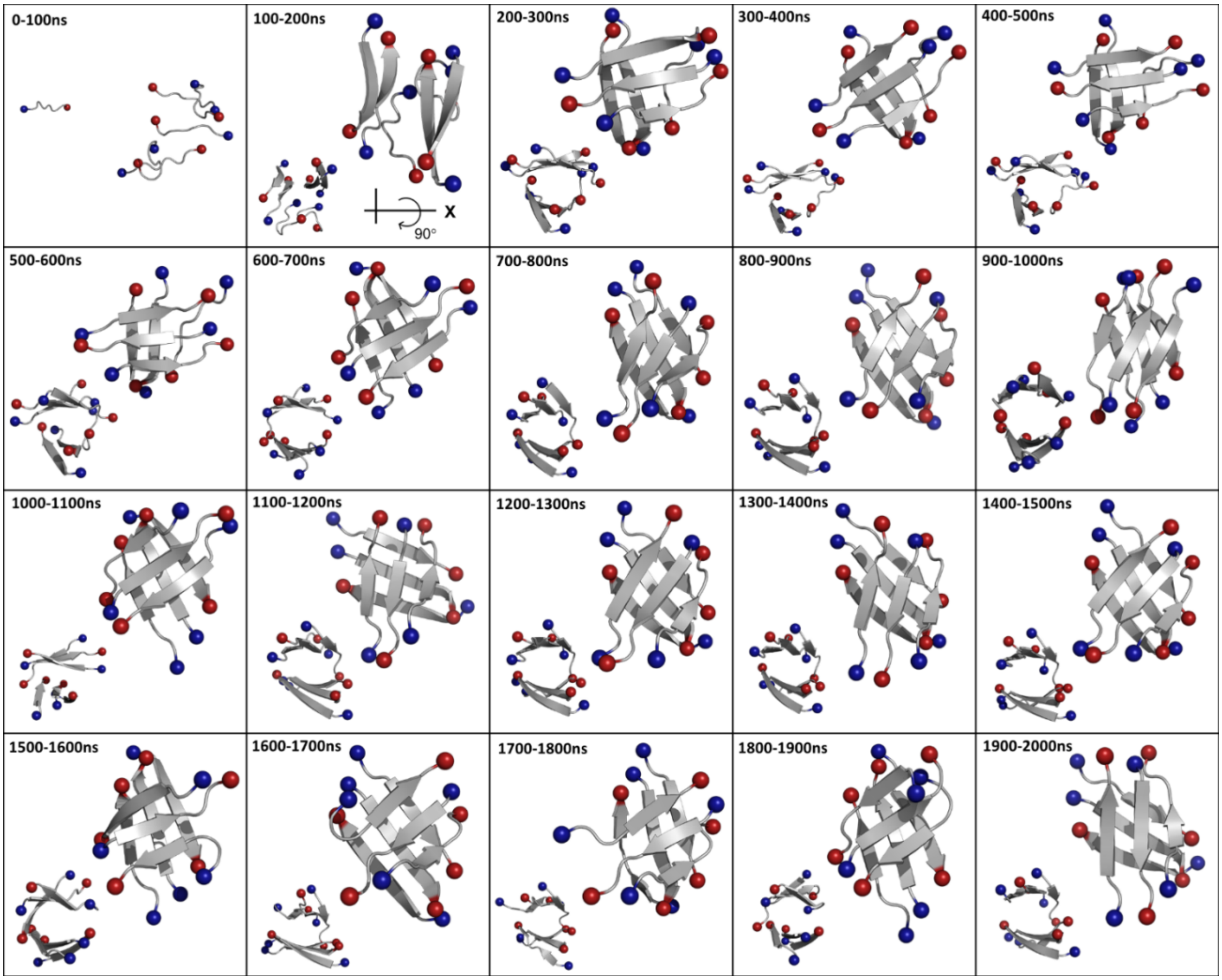


**Figure S8. Clustering analysis of Aβ$_{(16\text{-}22)}$ control replicate 1.** Dominant cluster morphology for each 100 ns interval of total 2 μs simulation time. Structures in the top right corner of each box are top-down views of the system, while the smaller structures in the bottom left corners show a side view of the system.

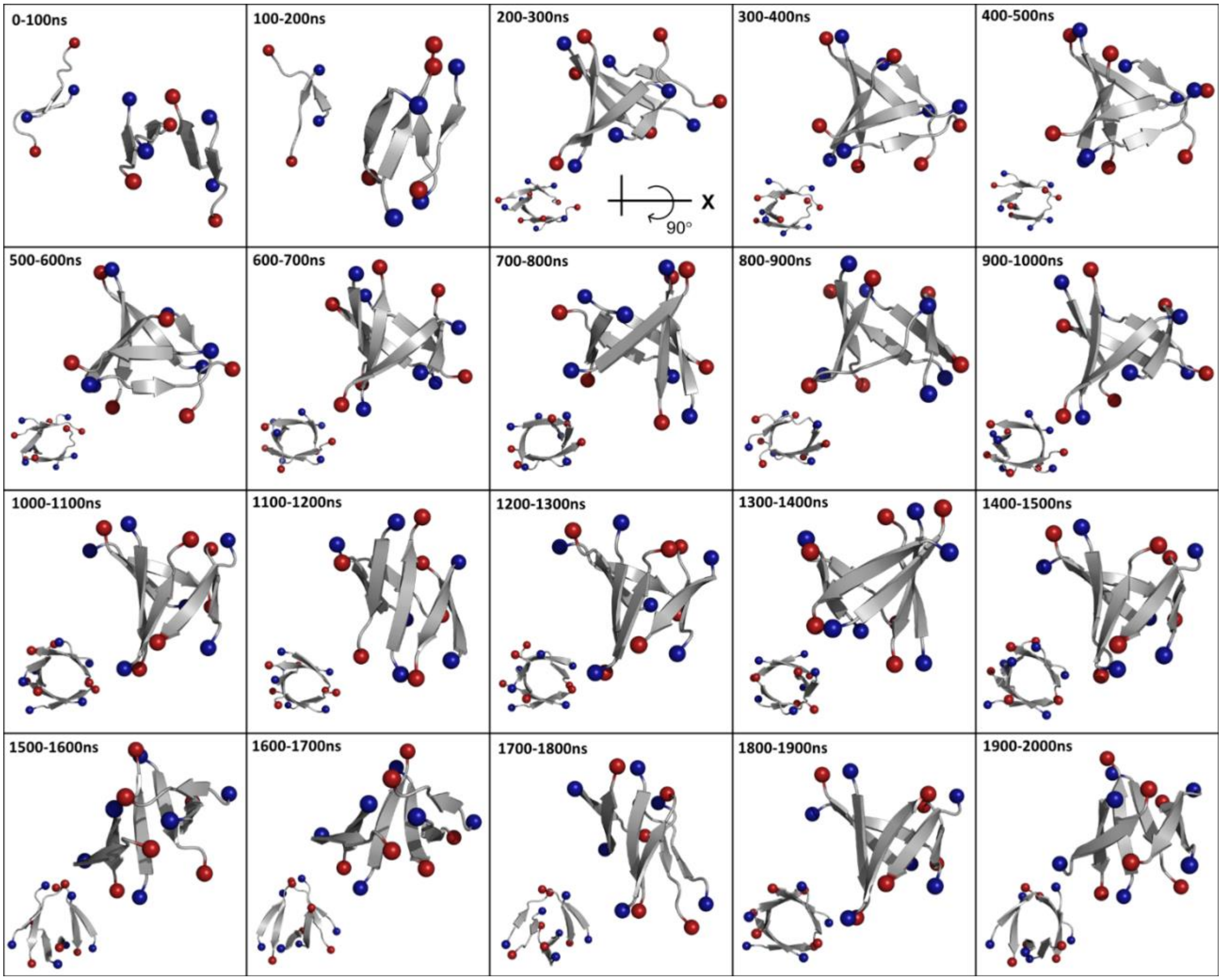


**Figure S9. Clustering analysis of Aβ$_{(16\text{-}22)}$ control replicate 2.** Dominant cluster morphology for each 100 ns interval of total 2 μs simulation time. Structures in the top right corner of each box are top-down views of the system, while the smaller structures in the bottom left corners show a side view of the system.

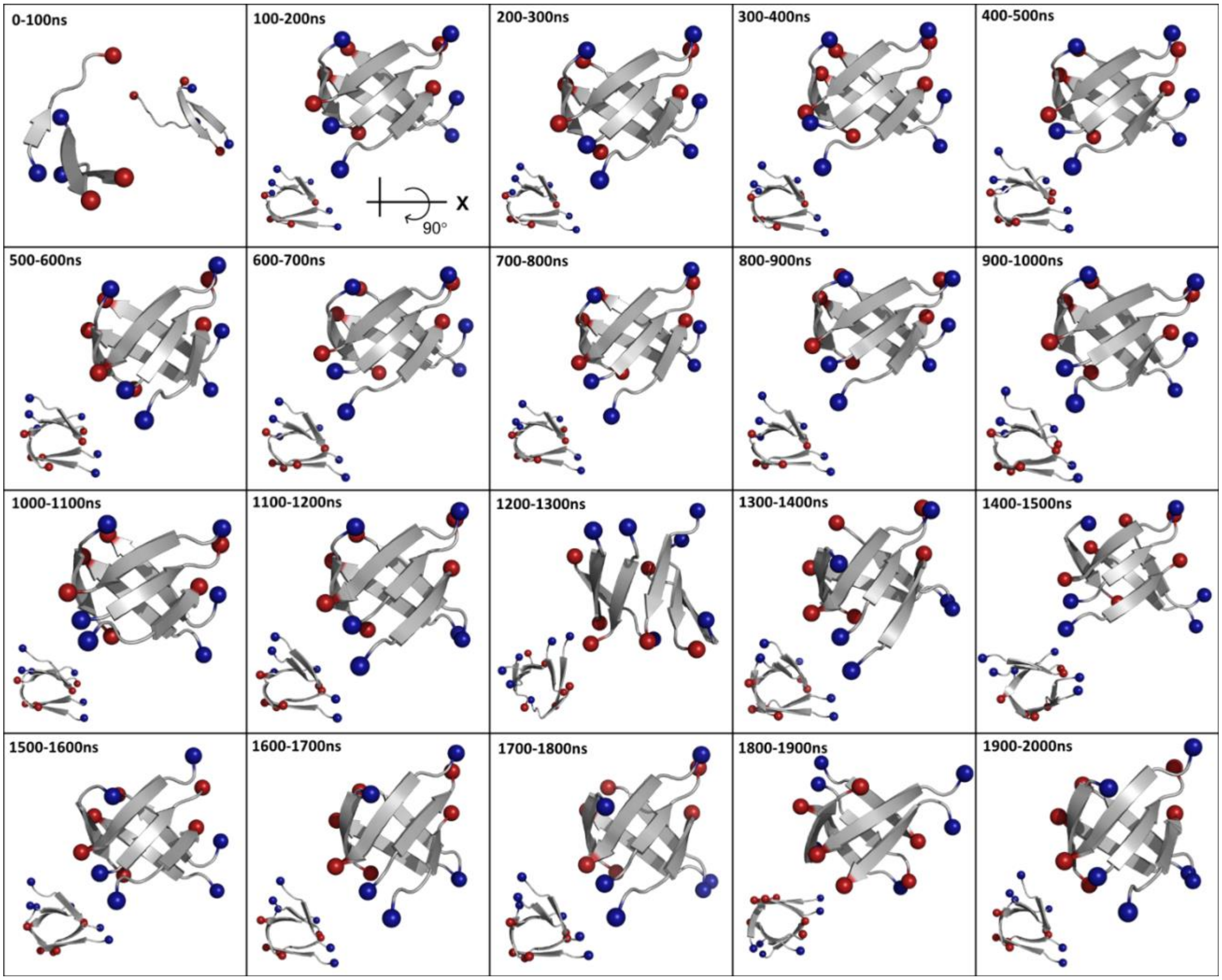


**Figure S10. Clustering analysis of A$\beta_{(16\text{-}22)}$ control replicate 3.** Dominant cluster morphology for each 100 ns interval of total 2 μs simulation time. Structures in the top right corner of each box are top-down views of the system, while the smaller structures in the bottom left corners show a side view of the system.

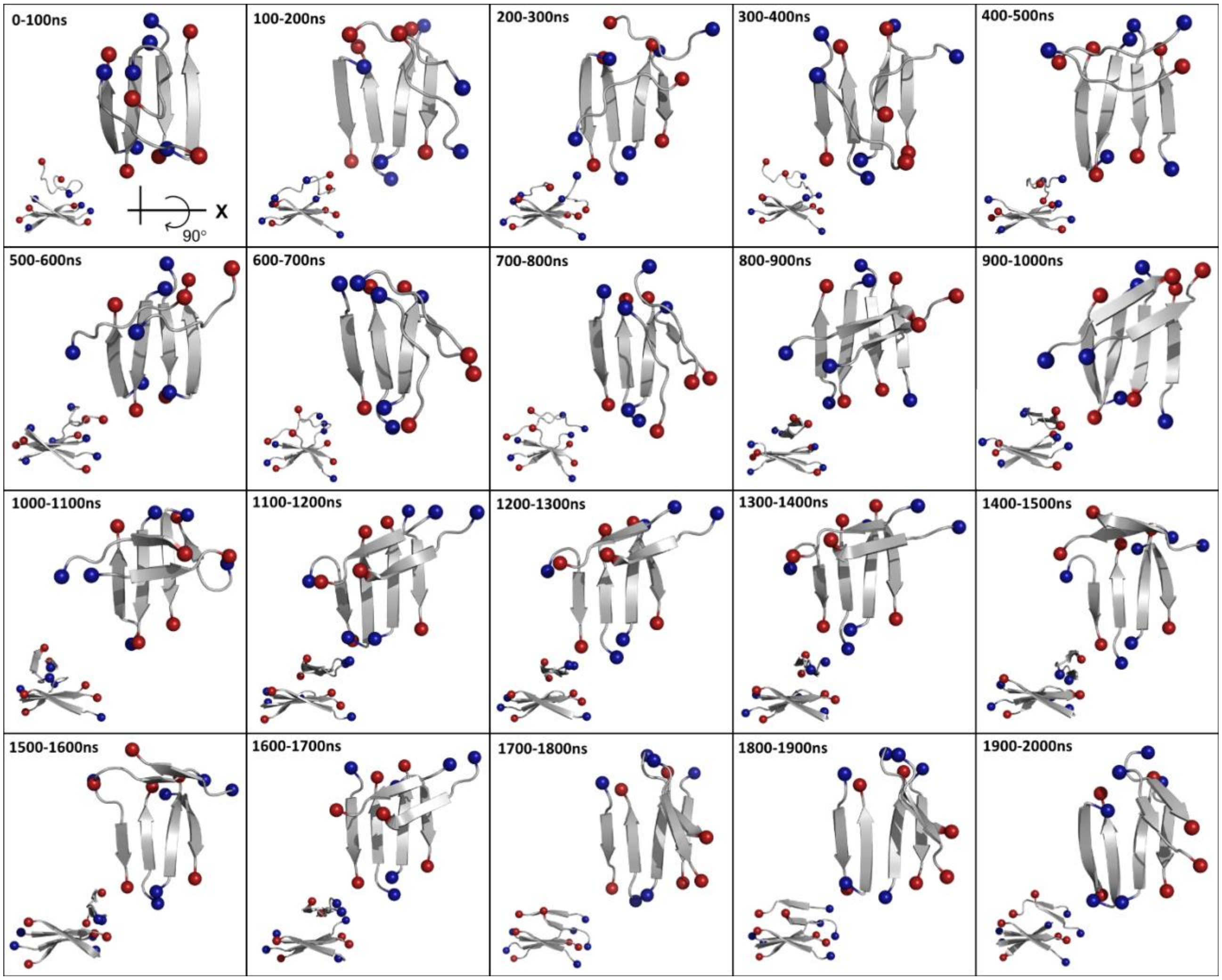


**Figure S11. Clustering analysis of Aβ(16-22) control replicate 4.** Dominant cluster morphology for each 100 ns interval of total 2 μs simulation time. Structures in the top right corner of each box are top-down views of the system, while the smaller structures in the bottom left corners show a side view of the system.

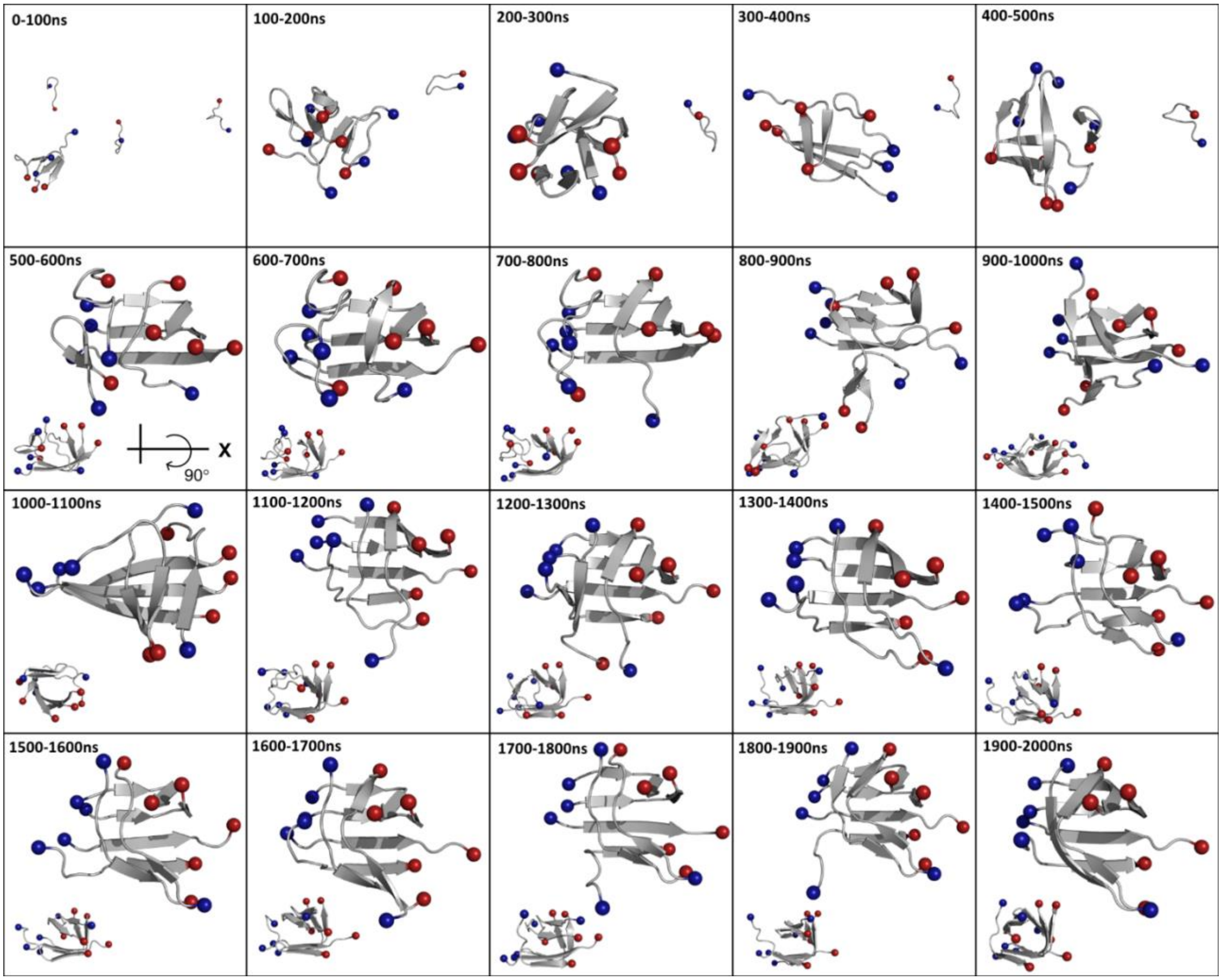


**Figure S12. Clustering analysis of IAPP$_{(20\text{-}29)}$ control replicate 1.** Dominant cluster morphology for each 100 ns interval of total 2 μs simulation time. Structures in the top right corner of each box are top-down views of the system, while the smaller structures in the bottom left corners show a side view of the system.

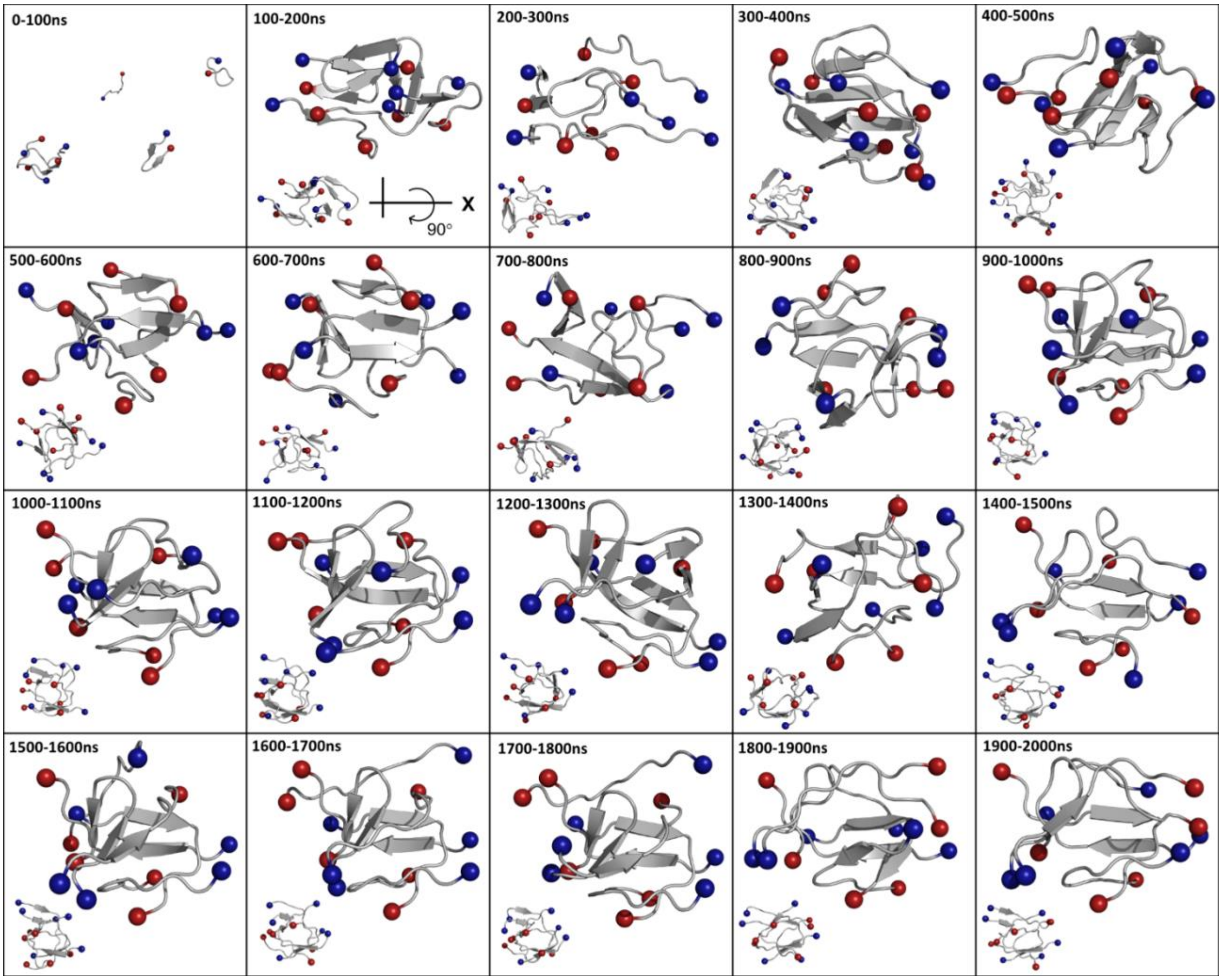


**Figure S13. Clustering analysis of IAPP$_{(20-29)}$ control replicate 2.** Dominant cluster morphology for each 100 ns interval of total 2 μs simulation time. Structures in the top right corner of each box are top-down views of the system, while the smaller structures in the bottom left corners show a side view of the system.

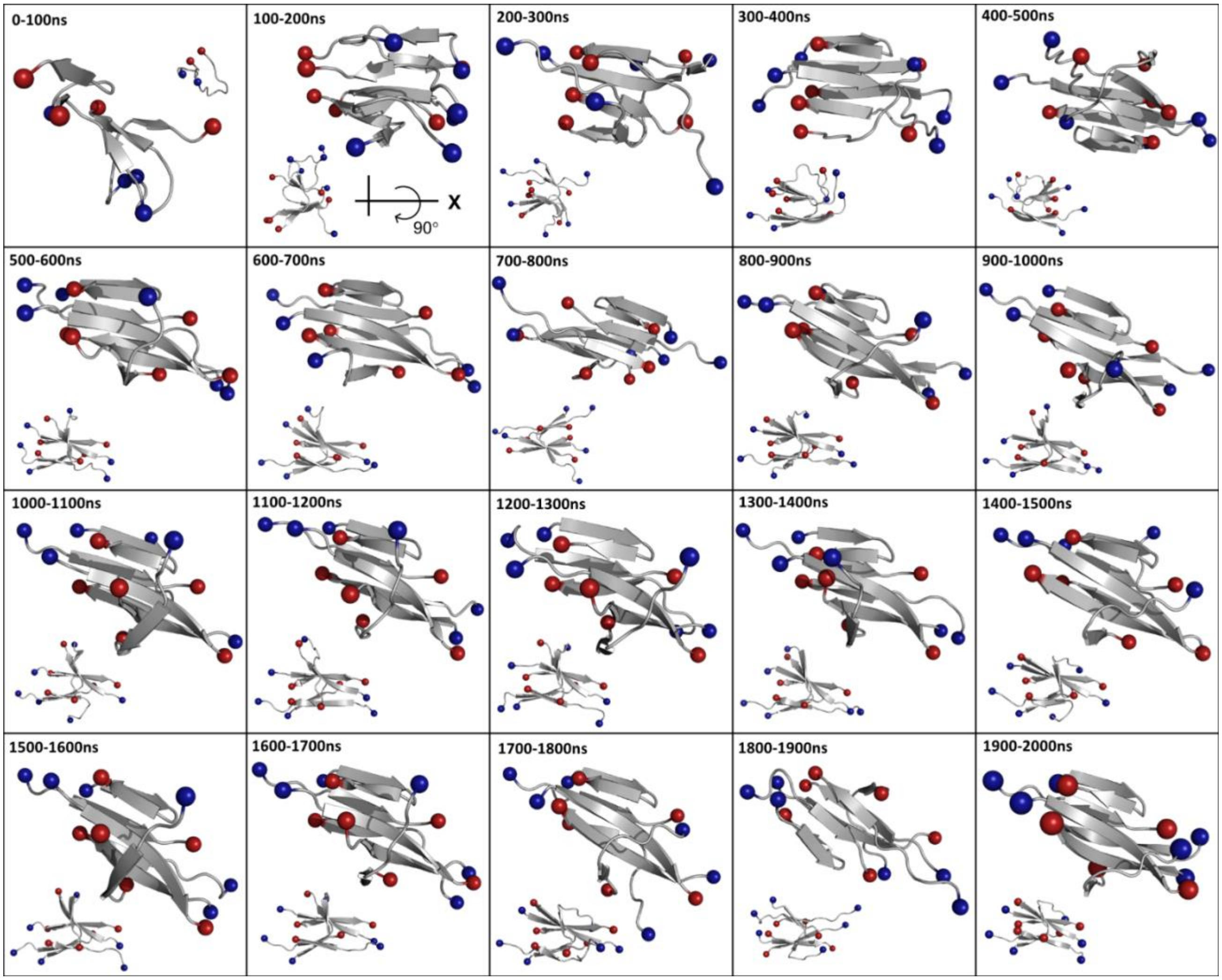


**Figure S14. Clustering analysis of IAPP$_{(20\text{-}29)}$ control replicate 3.** Dominant cluster morphology for each 100 ns interval of total 2 μs simulation time. Structures in the top right corner of each box are top-down views of the system, while the smaller structures in the bottom left corners show a side view of the system.

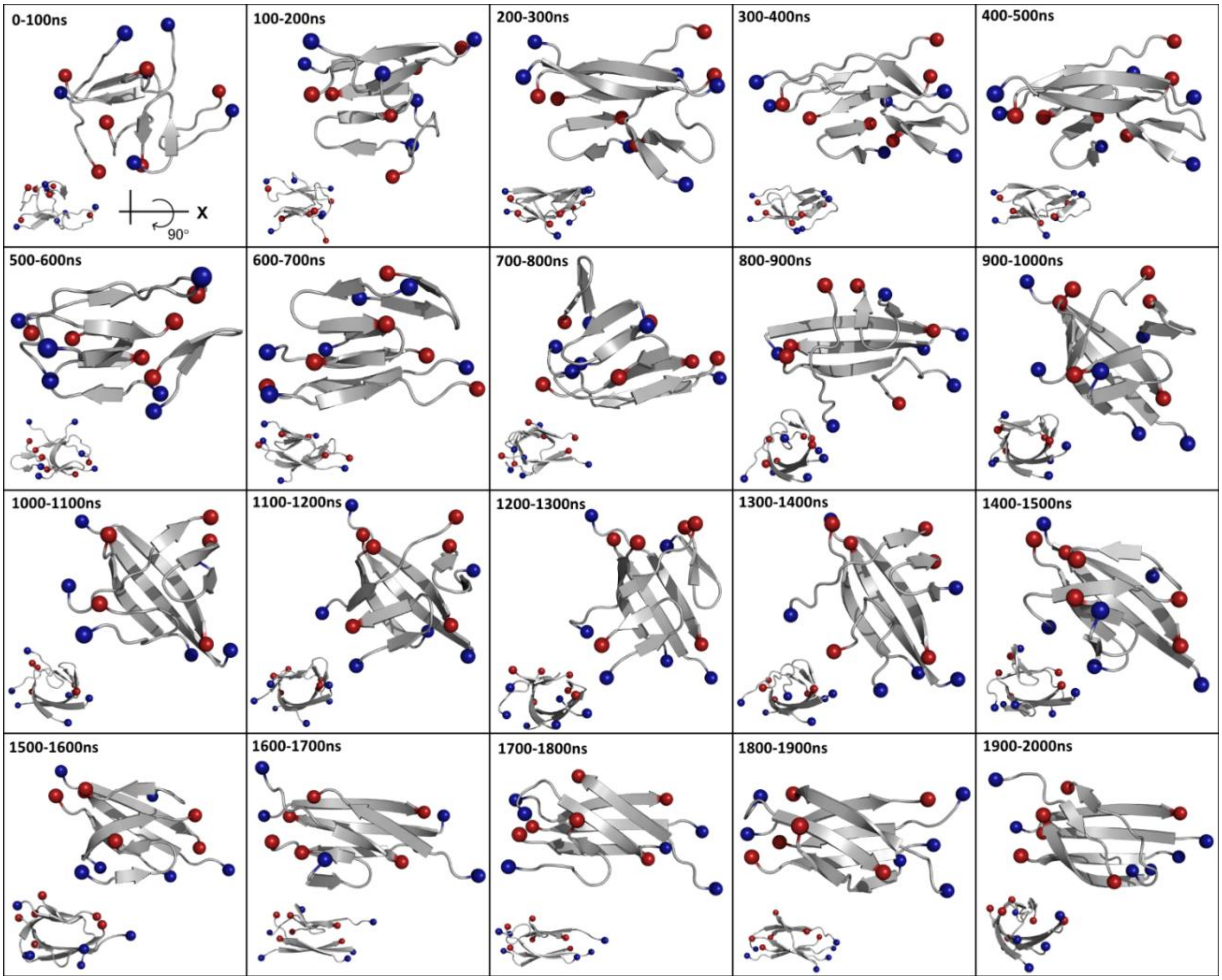


**Figure S15. Clustering analysis of IAPP$_{(20\text{-}29)}$ control replicate 4.** Dominant cluster morphology for each 100 ns interval of total 2 μs simulation time. Structures in the top right corner of each box are top-down views of the system, while the smaller structures in the bottom left corners show a side view of the system.

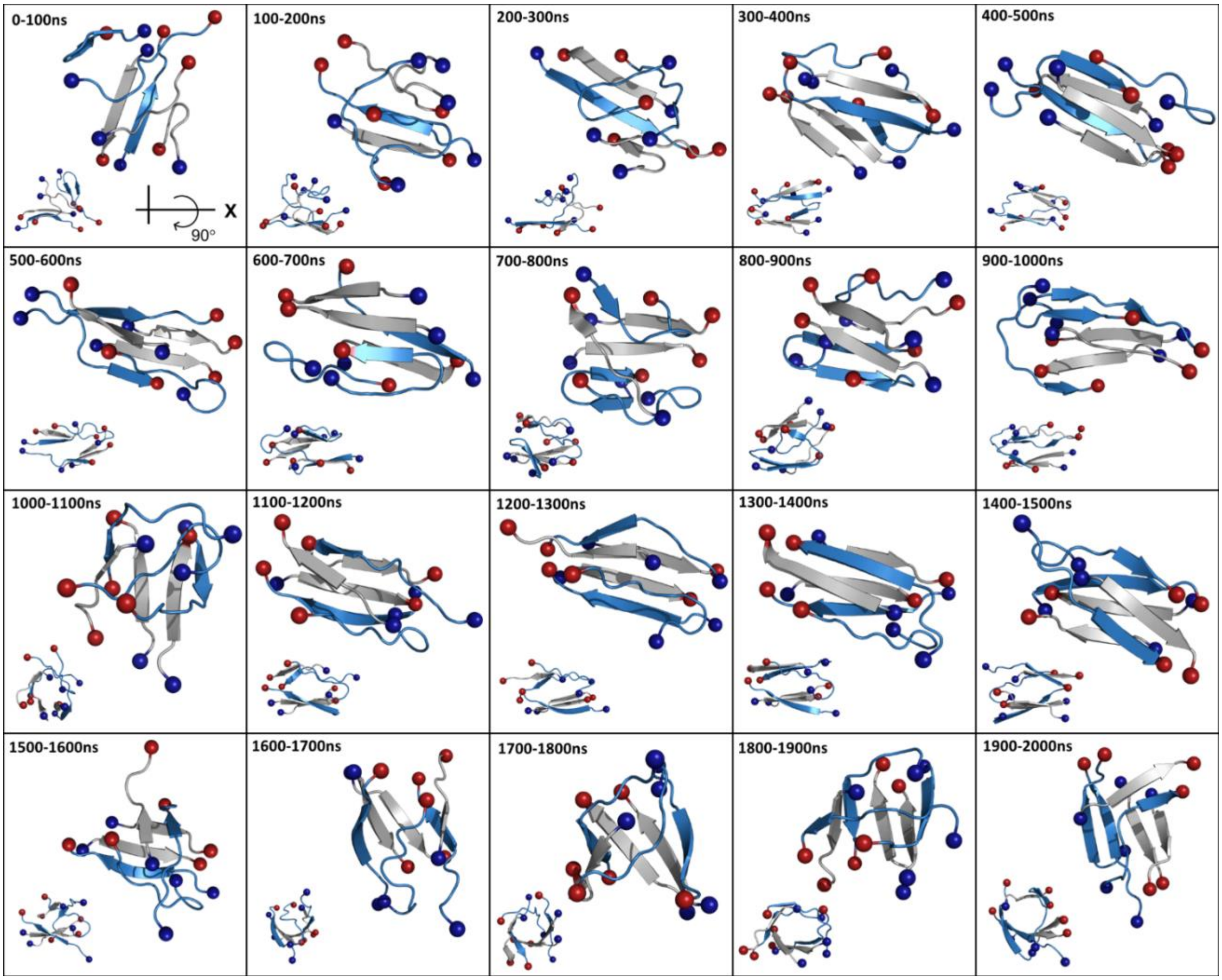


**Figure S16. Clustering analysis of Aβ$_{(16\text{-}22)}$+IAPP$_{(20\text{-}29)}$ replicate 1.** Dominant cluster morphology for each 100 ns interval of total 2 μs simulation time. Structures in the top right corner of each box are top-down views of the system, while the smaller structures in the bottom left corners show a side view of the system. IAPP$_{(20\text{-}29)}$ fragments are shown in blue, while Aβ$_{(16\text{-}22)}$ fragments are shown in gray.

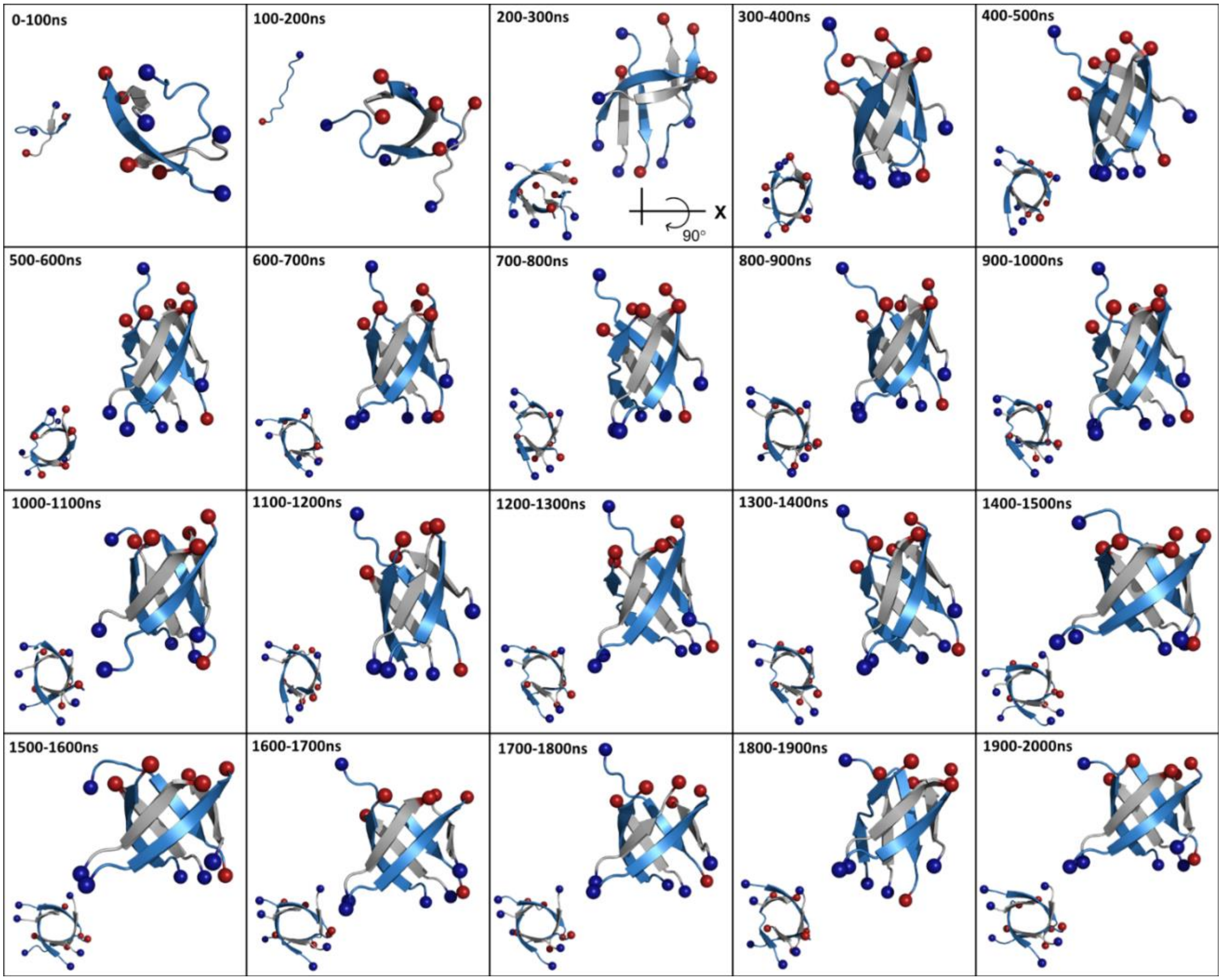


**Figure S17. Clustering analysis of Aβ$_{(16\text{-}22)}$+IAPP$_{(20\text{-}29)}$ replicate 2.** Dominant cluster morphology for each 100 ns interval of total 2 μs simulation time. Structures in the top right corner of each box are top-down views of the system, while the smaller structures in the bottom left corners show a side view of the system. IAPP$_{(20\text{-}29)}$ fragments are shown in blue, while Aβ$_{(16\text{-}22)}$ fragments are shown in gray.

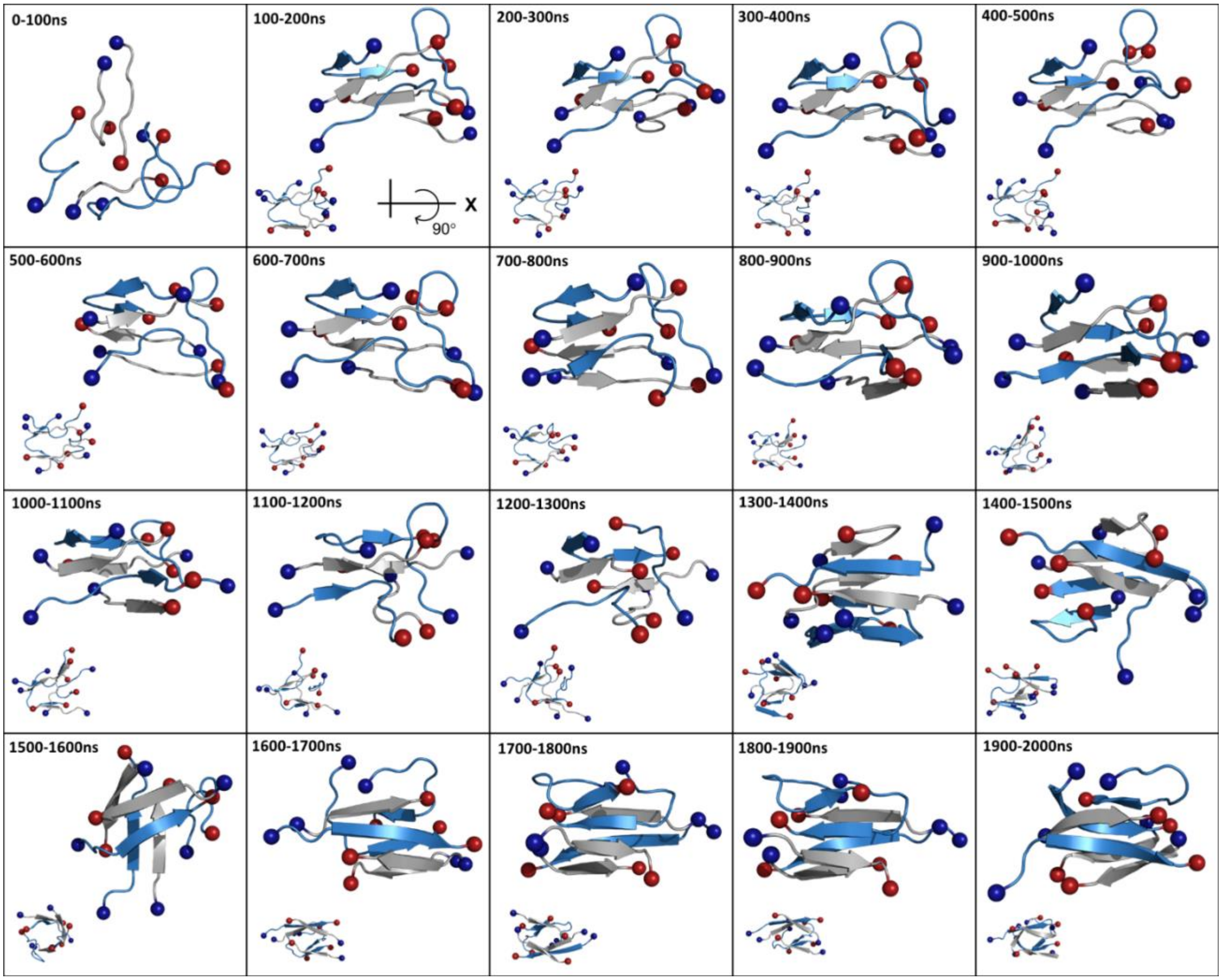


**Figure S18. Clustering analysis of Aβ$_{(16\text{-}22)}$+IAPP$_{(20\text{-}29)}$ replicate 3.** Dominant cluster morphology for each 100 ns interval of total 2 μs simulation time. Structures in the top right corner of each box are top-down views of the system, while the smaller structures in the bottom left corners show a side view of the system. IAPP$_{(20\text{-}29)}$ fragments are shown in blue, while Aβ$_{(16\text{-}22)}$ fragments are shown in gray.

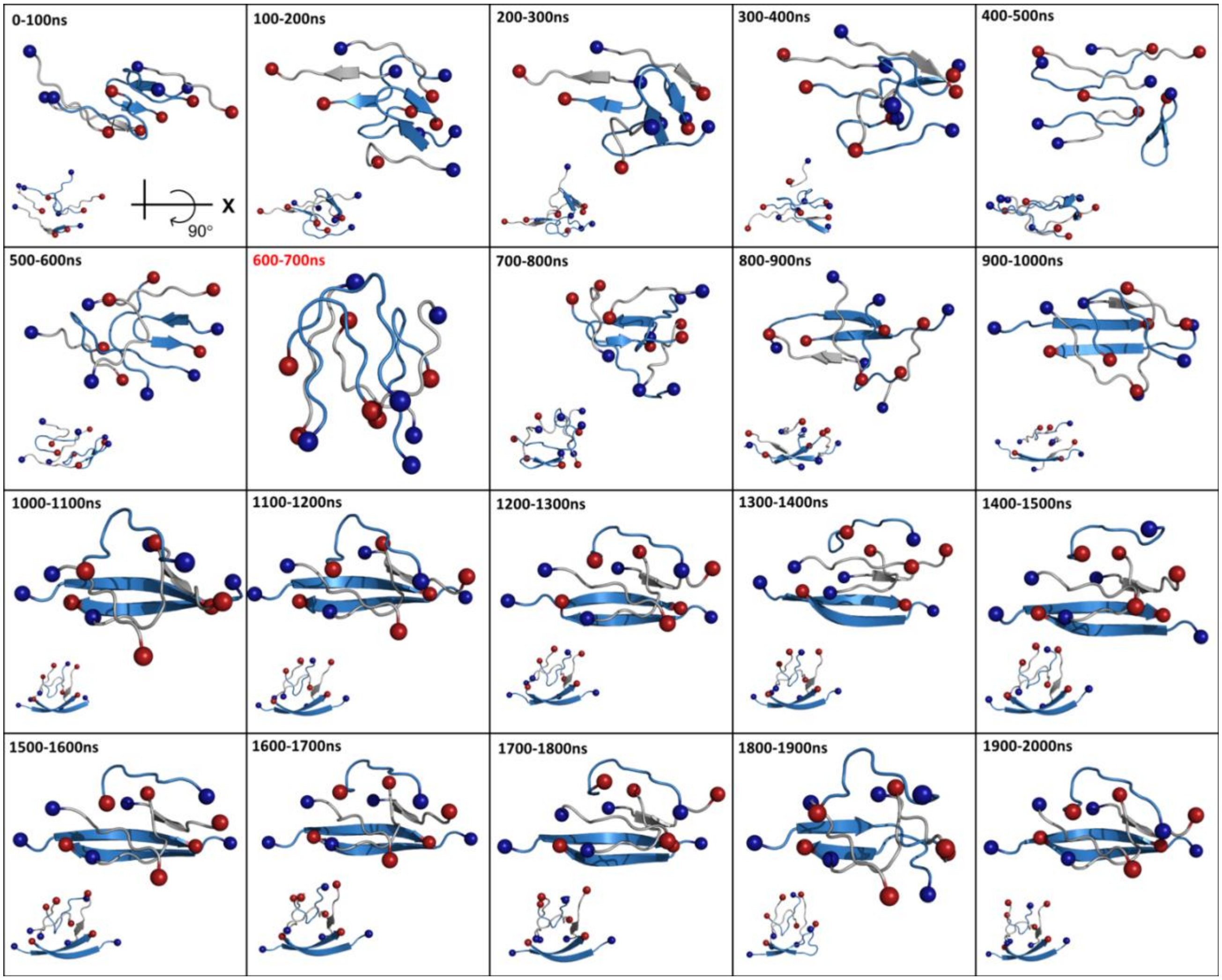


**Figure S19. Clustering analysis of Aβ(16-22)+IAPP(20-29) replicate 4.** Dominant cluster morphology for each 100 ns interval of total 2 μs simulation time. Structures in the top right corner of each box are top-down views of the system, while the smaller structures in the bottom left corners show a side view of the system. IAPP(20-29) fragments are shown in blue, while Aβ(16-22) fragments are shown in gray.

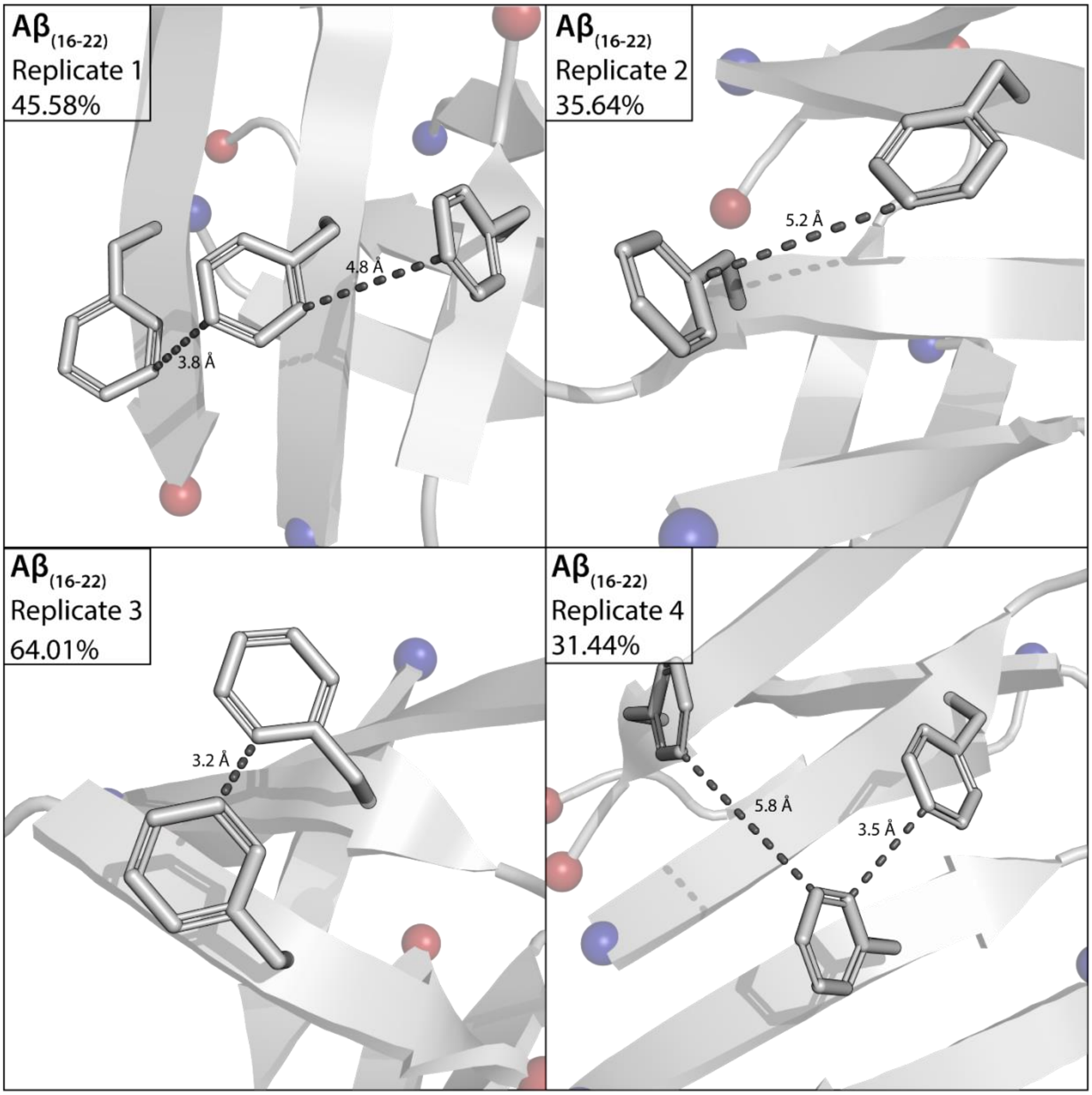


**Figure S20. Solvent-accessible phenylalanine interactions from RMSD cluster structures for Aβ$_{(16-22)}$ replicates.** Phenylalanine residues are shown as sticks, and colored gray. Peptides are shown as cartoon, and colored gray. The N- and C- termini are shown as spheres, and colored blue and red, respectively. Distance measurements are given in Angstroms. Structures shown are the dominant cluster morphology obtained from backbone RMSD clustering over the final 500 ns of each simulation. Percentages indicate the percentage of frames belonging to the most dominant cluster over the last 500 ns of simulation time.

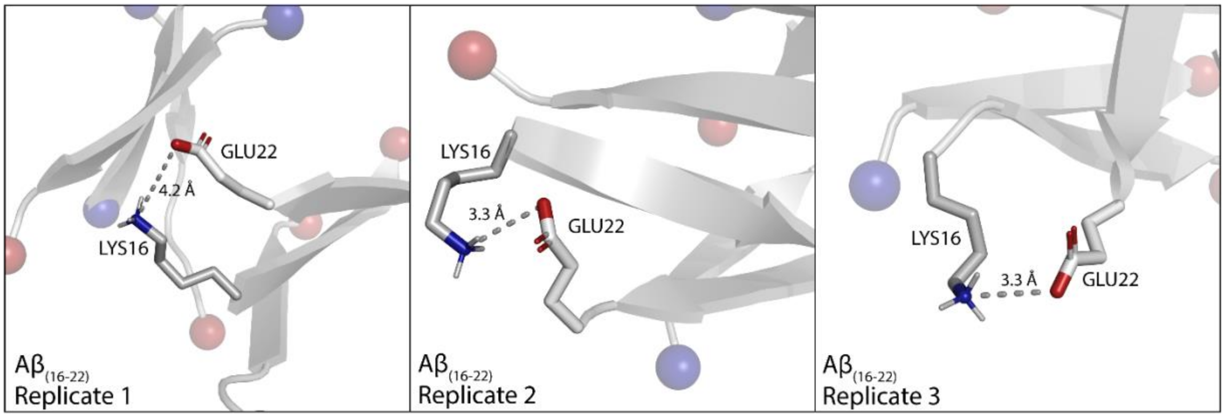


**Figure S21. Potential salt bridge interactions from Aβ$_{(16\text{-}22)}$ replicates.** Lysine and glutamate residues are shown as sticks, colored by element (gray carbon, blue nitrogen, red oxygen). Peptides are shown as cartoon, and colored gray. The N- and C- termini are shown as spheres, and colored blue and red, respectively. Distance measurements are given in Angstroms. Structures shown are the dominant cluster morphology obtained from backbone RMSD clustering over the final 500 ns of each simulation. Percentages indicate the percentage of frames belonging to the most dominant cluster over the last 500 ns of simulation time.

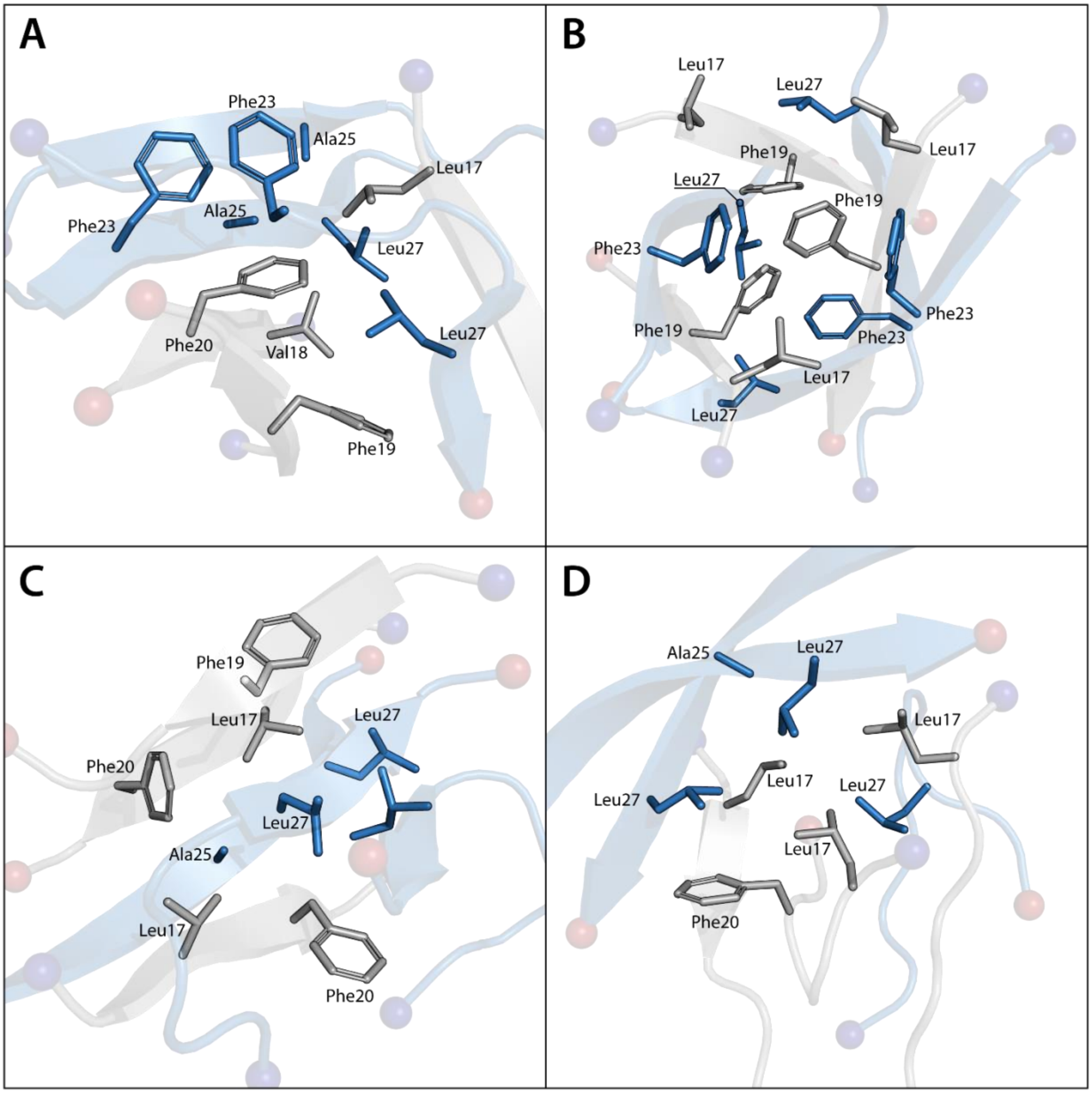


**Figure S22. Hydrophobic packing in heterogeneous Aβ$_{(16\text{-}22)}$ + IAPP$_{(20\text{-}29)}$ systems.** Residues involved in hydrophobic packing are shown as sticks. Aβ$_{(16\text{-}22)}$ is colored gray, and IAPP$_{(20\text{-}29)}$ is colored blue. The N- and C- termini are shown as spheres, and colored blue and red, respectively. Distance measurements are given in Angstroms. Structures shown are the dominant cluster morphology obtained from backbone RMSD clustering over the final 500 ns of each simulation. Percentages indicate the percentage of frames belonging to the most dominant cluster over the last 500 ns of simulation time.

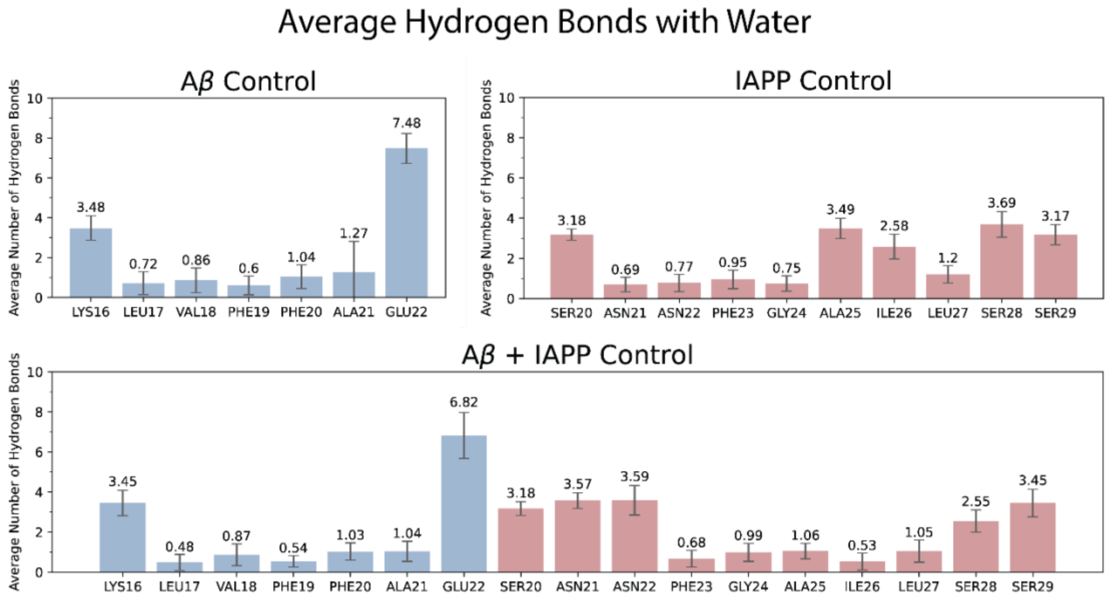


**Figure S23. Average hydrogen bonds per residue with water.** Averages represent the average across all replicates over the last 500 ns of simulation.

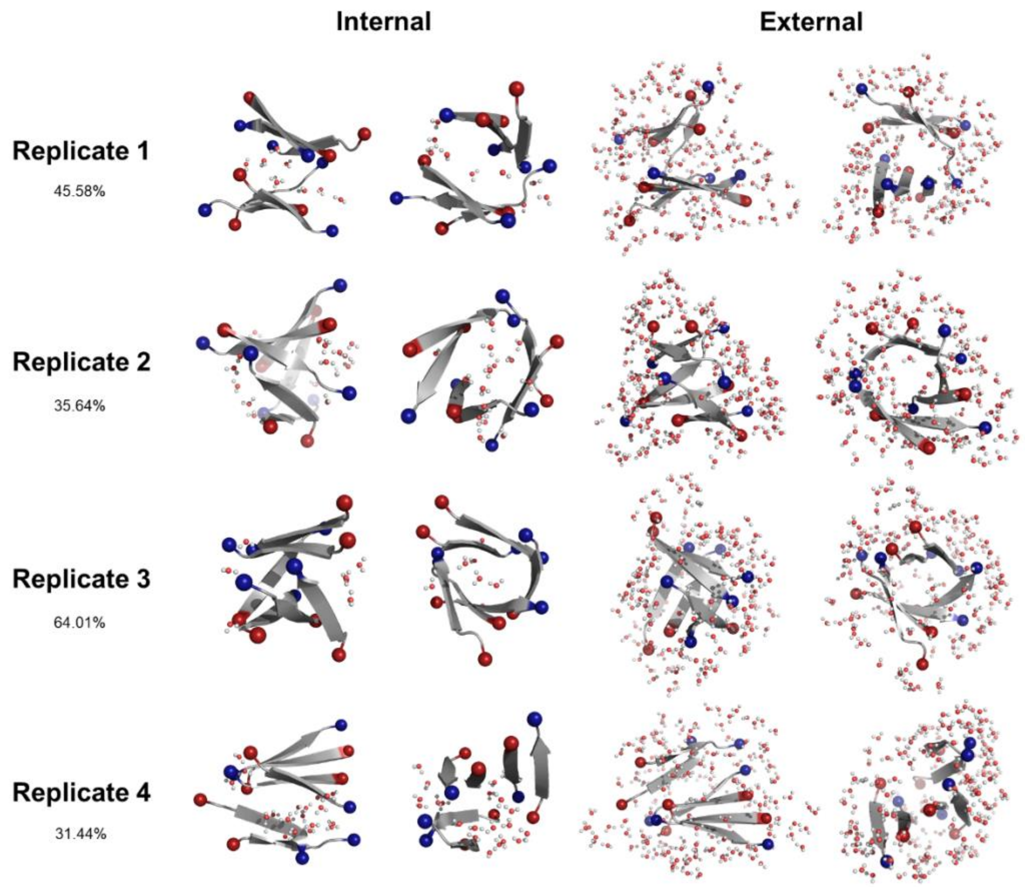


**Figure S24. Internal and external hydration shells of dominant Aβ control hexamer morphologies.** Each morphology represents the central structure of the dominant cluster obtained from the last 0.5 μs of each system. Leftmost structures in each column are top-down views of the hexamers, while right side structures are side views meant to show the internal cavity of barrel-like structures. Water contacts were defined as external if they were within 3 angstroms of surface peptide residues. Water contacts were defined as internal if their distance from the center of mass of the hexamer was less than the distance of most peptide residues from the center of mass of the hexamer.

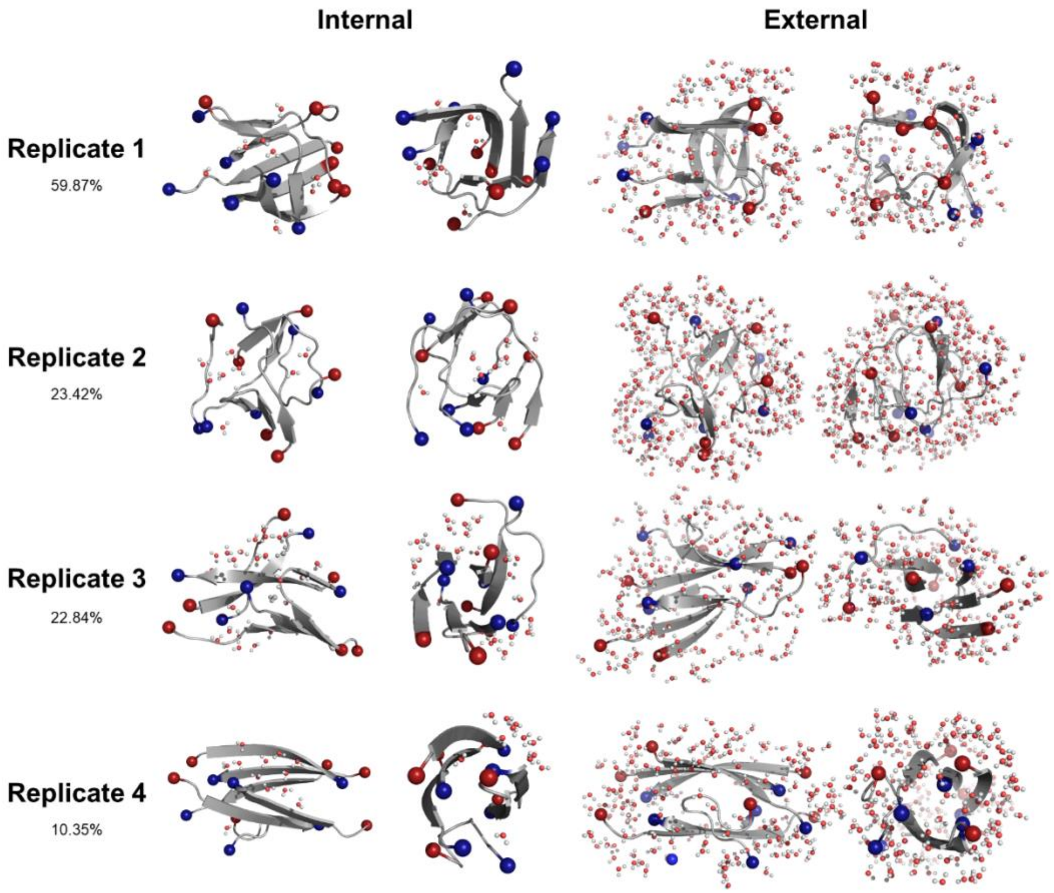


**Figure S25. Internal and external hydration shells of dominant IAPP control hexamer morphologies.** Each morphology represents the central structure of the dominant cluster obtained from the last 0.5 µs of each system. Leftmost structures in each column are top-down views of the hexamers, while right side structures are side views meant to show the internal cavity of barrel-like structures. Water contacts were defined as external if they were within 3 angstroms of surface peptide residues. Water contacts were defined as internal if their distance from the center of mass of the hexamer was less than the distance of most peptide residues from the center of mass of the hexamer.

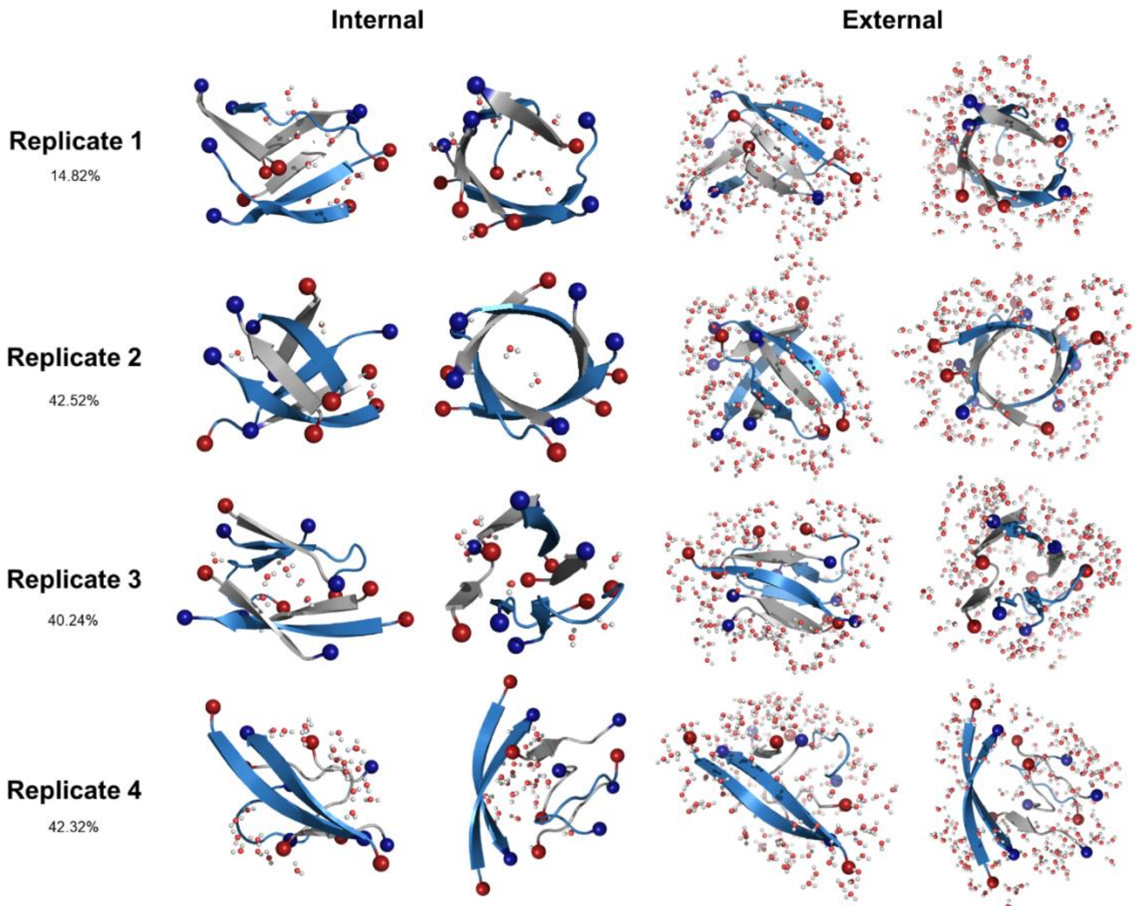


**Figure S26. Internal and external hydration shells of dominant Aβ+IAPP hexamer morphologies.** Each morphology represents the central structure of the dominant cluster obtained from the last 0.5 µs of each system. $IAPP_{(20-29)}$ fragments are shown in blue, while $A\beta_{(16-22)}$ fragments are shown in gray. Leftmost structures in each column are top-down views of the hexamers, while right side structures in each column are side views intended to show the internal cavity of barrel-like structures. Water contacts were defined as external if they were within 3 angstroms of surface peptide residues. Water contacts were defined as internal if their distance from the center of mass of the hexamer was less than the distance of most peptide residues from the center of mass of the hexamer.
